# stAPAminer: Mining Spatial Patterns of Alternative Polyadenylation for Spatially Resolved Transcriptomic Studies

**DOI:** 10.1101/2022.07.20.500789

**Authors:** Guoli Ji, Qi Tang, Sheng Zhu, Junyi Zhu, Pengchao Ye, Shuting Xia, Xiaohui Wu

**Affiliations:** Pasteurien College, Soochow University, Suzhou 215000, China; Department of Automation, Xiamen University, Xiamen 361005, China; Institute of Neuroscience, Soochow University, Suzhou 215000, China

**Keywords:** Alternative polyadenylation, Spatial transcriptomics, Single-cell RNA sequencing, Spatial pattern, Imputation

## Abstract

Alternative polyadenylation (APA) contributes to transcriptome complexity and gene expression regulation, which has been implicated in various cellular processes and diseases. Single-cell RNA-seq (scRNA-seq) has led to the profile of APA at the single-cell level, however, the spatial information of cells is not preserved in scRNA-seq. Alternatively, spatial transcriptomics (ST) technologies provide opportunities to decipher the spatial context of the transcriptomic landscape within single cells and/or across tissue sections. Pioneering studies on ST have unveiled potential spatially variable genes and/or splice isoforms, however, the pattern of APA usages in spatial contexts remains unappreciated. Here, we developed a toolkit called stAPAminer for mining spatial patterns of APA from spatial barcoded ST data. APA sites were identified and quantified from the ST data. Particularly, an imputation model based on K-nearest neighbors algorithm was designed for recovering APA signals. Then APA genes with spatial patterns of APA usage variation were identified. By analyzing the well-established ST data of mouse olfactory bulb (MOB), we present a detailed view of spatial APA usage across morphological layers of MOB with stAPAminer. We complied a comprehensive list of genes with spatial APA dynamics and obtained several major spatial expression patterns representing spatial APA dynamics in different morphological layers. Extending this analysis to two additional replicates of the MOB ST data, we found that spatial APA patterns of many genes are reproducible among replicates. stAPAminer employs the power of ST for exploring transcriptional atlas of spatial APA patterns with spatial resolution, which is available at https://github.com/BMILAB/stAPAminer.

## Introduction

mRNA polyadenylation is a critical mRNA processing event occurring towards the completion of transcription, which involves two tightly coupled steps: cleavage at the nascent transcript followed by the addition of an untemplated poly(A) tail to its 3′ end [1, 2]. Over 70% of mammalian genes possess more than one poly(A) site, providing the possibility of the modulated use of selective poly(A) sites through alternative polyadenylation (APA) [1, 2]. APA contributes greatly to the complexity of transcriptome and proteome by generating isoforms of the same gene with distinct 3′ UTRs or coding regions. APA is dynamically regulated in various cellular processes and diseases, such as cell activation, proliferation differentiation, neurodegenerative disorders, and cancer [3–12]. Studies using bulk 3′-end sequencing (reviewed in [2, 13, 14]) and/or RNA-seq (reviewed in [14, 15]) have revealed extensive 3′ UTR lengthening/shortening events in various processes. For example, 3′ UTRs generally shorten in proliferating cells, while 3′ UTRs lengthen during embryonic differentiation [16] and animal neurogenesis [17, 18].

In contrast to bulk methodologies, single-cell RNA sequencing (scRNA-seq) protocols characterize transcriptional landscape in individual cells, with many protocols utilizing 3′ selection/enrichment in library construction, such as Drop-seq [19], CEL-seq [20], and 10x Genomics [21]. Although most scRNA-seq studies initially focus only on gene expression profiling, these scRNA-seq technologies inherently capture a surprising amount of information about isoform usage, providing great potentials to profile APA events at the single-cell level. Computational tools, such as scDAPA [22], scAPA [23], Sierra [24], scAPAtrap [25], and scDaPars [26], have been proposed to identify APA sites in single cells and/or profile differential APA isoform usage among cell types from scRNA-seq. Particularly, single-cell APA profile compiled from scRNA-seq enables the discovery of hidden subpopulations of cells that are unrecognizable in conventional gene expression analysis [25, 26], revealing the possibility of discerning cell identities with the APA layer independent of gene expression.

While scRNA-seq is powerful in profiling transcriptome in individual cells, the spatial information of cells is not preserved due to the tissue dissociation before sequencing. Characterizing the spatial organization and molecular features of cells is essential to understand cellular interactions and organization in the tissue microenvironment. Several strategies for spatial transcriptomics (ST) have been established for measuring spatially resolved gene expression [27], such as MERFISH [28], seqFISH [29], and spatial transcriptomics through spatial barcoding [30], providing opportunities to decipher the spatial context of the transcriptomic landscape within single cells and/or across tissue sections. The identification of spatially variable genes is usually the critical first step in analyzing ST data to spatially resolve the transcriptomic landscape across tissues. Recently, a few computational approaches have been proposed for exploring spatial gene expression trends, including SpatialDE [31], Trendsceek [32], SPARK [33], and SPARK-X [34]. Continuous gradients or spatial gene expression patterns can be identified by these tools, which contributes to disclosing significant biological discoveries that otherwise cannot be revealed by scRNA-seq alone. Although most spatially resolved transcriptomic studies have restricted analysis at the gene level, these studies, especially the ST method of spatial barcoding, may provide additional information on transcript isoforms, enabling multiple layers of transcriptome information to be obtained from ST experiments without changing experimental methods. Lately, dynamic alternative splicing and brain-region-specific isoform expression have been observed using a single-cell isoform RNA sequencing technology [35]. These pioneering studies have implicated potential spatially variable genes and/or splice isoforms, however, the pattern of APA usages in spatial contexts remains unappreciated.

Here, we developed a toolkit called stAPAminer for mining spatial patterns of APA from spatial barcoded ST data. First, poly(A) sites were identified from ST data and then APA site usages of genes in individual spots were quantified. Particularly, an imputation model based on K-nearest neighbors algorithm was designed for recovering APA signals obtained from the ST data by borrowing information from the spatial gene expression profile, which can effectively mitigate the noise and bias caused by the dropout phenomenon. Based on the profile of imputed APA usages, APA genes with differential APA usage between morphological layers and genes with spatial patterns of APA usage variation were identified. By analyzing the well-established ST data of mouse olfactory bulb (MOB) data, we present a detailed view of spatial APA usage across MOB regions with stAPAminer. We complied a comprehensive list of genes with spatial APA dynamics and obtained several major spatial APA patterns representing APA dynamics in different morphological layers. These genes are enriched in gene ontology terms directly associated with olfactory bulb development, highlighting the benefits of spatial APA analysis with scAPAminer. Extending the analysis to two additional replicates of the MOB ST data, we found spatial APA patterns of many genes are reproducible among replicates, demonstrating the robustness and effectiveness of scAPAminer in spatial APA analysis. stAPAminer employs the power of ST for exploring transcriptional atlas of spatial APA patterns with spatial resolution variation, establishing an additional layer of gene expression at the isoform resolution.

## Results

### stAPAminer was developed for the analysis of spatial APA dynamics from ST data

We developed an R package, stAPAminer, for exploring spatial APA dynamics from ST data (**Figure 1**). First poly(A) sites are identified and quantified from ST data by existing tools like scAPAtrap used in this study (Figure 1A). Then a poly(A) site expression matrix is obtained which is similar to conventional gene-cell expression matrix obtained from scRNA-seq, except that each row denotes one poly(A) site rather than a gene and each column denotes a spot. Next, a matrix of 3′ UTR or intronic APA usage is obtained from the poly(A) site expression matrix (Figure 1B). For 3′ UTR APA analysis, genes with at least two 3′ UTR poly(A) sites are extracted and the APA usage of each gene is represented by the relative usage of distal poly(A) site (RUD). For intronic APA analysis, APA genes with at least one intronic poly(A) site are extracted and the APA usage of each gene is represented by ratio of the intronic site (Materials and methods). The APA usage matrix is even sparser than the gene-spot expression matrix, therefore, we proposed an imputation model based on k-nearest neighbors (KNN) for recovering APA signals in the APA usage matrix. This model leverages the gene expression profile to infer spot-spot distance and then imputes the sparse APA usage matrix by borrowing information of APA usage from neighboring spots. The *k* value is the most critical parameter in the KNN-based imputation model, which can be determined using a strategy based on a comprehensive index (Materials and methods). After imputation, benchmarking analyses were conducted to investigate the performance of the imputation model. The imputed APA usage matrix was used for exploring spatial APA dynamics (Figure 1C), including the detection of genes with spatially variable usages of APA (SVAPA), genes with differential APA usages between layers (DEAPA), and genes with layer-specific APA (LSAPA). These three types of APA genes reflect the spatial characteristics of APA from different aspects, establishing the full landscape of spatial APA dynamics. Moreover, representative spatial APA patterns can be further obtained by clustering these APA genes. stAPAminer was implemented as an open source R package, available at https://github.com/BMILAB/stAPAminer.

**Figure 1.**
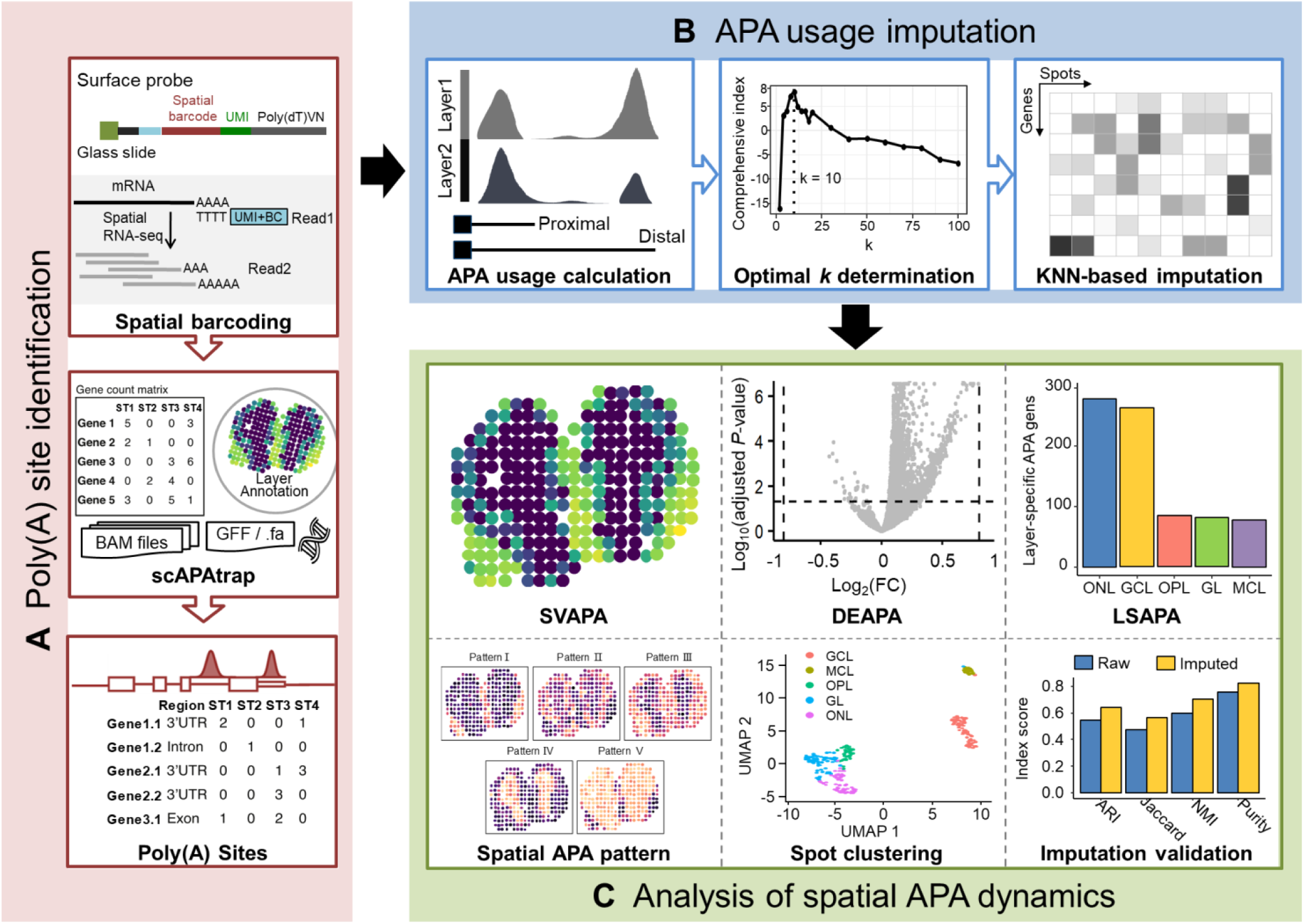
Schema of stAPAminer. **A.** Identification of poly(A) sites from spatial transcriptomics data using scAPAtrap. **B.** Quantification and imputation of APA usages. The sparse gene-spot matrix of APA usages is imputed by the KNN-based imputation model embedded in stAPAminer and the optimal *k* value is determined by using a comprehensive index. **C.** Analysis of spatial APA dynamics with stAPAminer. stAPAminer can be used to detect spatially variable APA (SVAPA), differential APA usages between layers (DEAPA), and layer-specific spatial APA events (LSAPA). APA, alternative polyadenylation; FC, fold change; KNN, k-nearest neighbors.

### Genome-wide poly(A) sites were identified from MOB ST data

We applied stAPAminer to analyze a spatial barcoded ST dataset of MOB [30]. Following other studies [31, 33], we mainly used the ‘MOB Replicate 11’ file for analysis, which consists of 11,274 genes on 260 spots after filtration (hereafter called ST-MOB). A total of 47,888 poly(A) sites were extracted using the scAPAtrap pipeline (**Figure 2A**), with an average of 14,436 poly(A) sites per spot (Table S1). Majority of poly(A) sites are located in 3′ UTRs/extended 3′ UTRs (21,671) or introns (17,645), which is consistent with the previous observation in scRNA-seq [24, 25]. The base composition surrounding 3′ UTR poly(A) sites from ST-MOB resembles the general profile of annotated poly(A) sites (Figure 2B). A high frequency of occurrence of core poly(A) hexamers is observed in the upstream poly(A) site region, including AAUAAA, AUUAAA, and GU-rich hexamers (Figure S1). Next, we used poly(A) sites identified from MOB scRNA-seq data (hereafter called SC-MOB) [36] and annotated poly(A) sites of bulk 3′-seq from PolyASite 2.0 [37] to validate poly(A) sites identified from ST-MOB. Up to 16,266 poly(A) sites and 28,278 poly(A) sites from ST-MOB were also found in SC-MOB and the PolyASite 2.0 database, respectively. While 24,161 poly(A) sites were exclusively found in ST-MOB, which may represent potential novel polyadenylation events in the ST data that cannot be detected from scRNA-seq or bulk 3′-seq. Poly(A) sites from ST-MOB are close to annotated poly(A) sites or sites from SC-MOB (Figure 2C). We noted that there is a sharp decrease at the zero position of SC-MOB, which may be due to the 1-bp bias of the position of poly(A) sites identified by scAPAtrap [25]. Since the expression of a poly(A) site is the read coverage of the peak region of a poly(A) site rather than a single position, the 1-bp bias would not affect the poly(A) site quantification and downstream analysis. Reassuringly, the poly(A) site expression profile of ST-MOB is highly correlated with that of SC-MOB (Pearson’s correlation =0.70, Figure 2D) and neural-related samples of bulk 3′-seq (Pearson’s correlation = 0.69, Figure S2). These observations demonstrated the authenticity of poly(A) sites identified from ST-MOB data.

**Figure 2.**
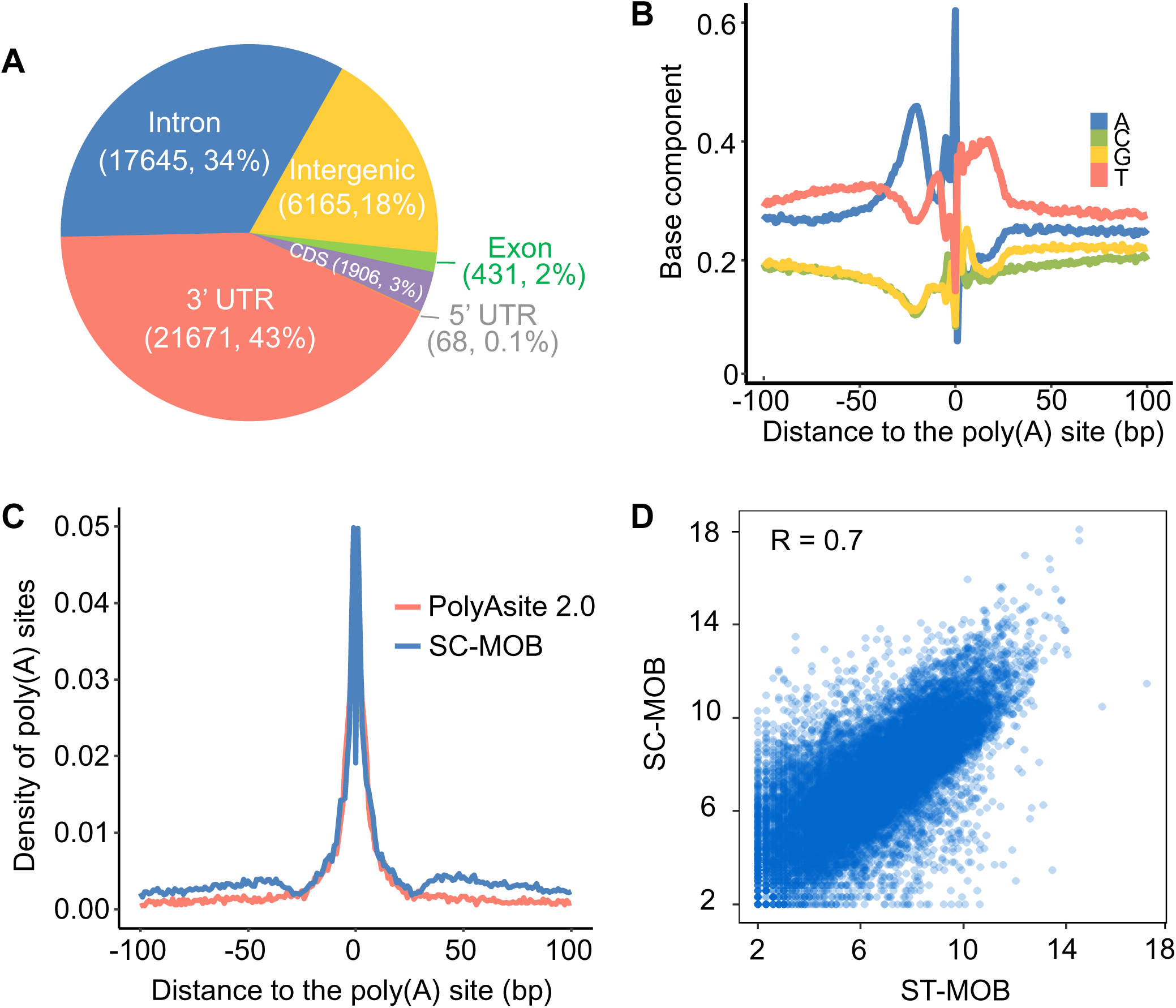
Validation of poly(A) sites identified from the ST-MOB data. **A.** Distribution of poly(A) sites from ST-MOB in different genomic regions. **B.** Nucleotide compositions of the sequences surrounding 3′ UTR poly(A) sites from ST-MOB. Y-axis denotes the fractional nucleotide content at each position. On the x-axis, “0” denotes the poly(A) site. **C.** Comparison of ST-MOB poly(A) sites with annotated poly(A) sites in PolyASite 2.0 database and poly(A) sites identified from scRNA-seq data of MOB (SC-MOB). The curves show the distance from ST-MOB poly(A) sites to annotated poly(A) sites and SC-MOB poly(A) sites. **D.** Scatter plot showing the correlation of poly(A) site expression profiles obtained from ST-MOB and SC-MOB. Each dot is one poly(A) site and the axis is log_2_ scaled. The Pearson’s correlation is indicated in the figure.

### stAPAminer effectively recovers highly sparse APA signals obtained from ST-MOB

After identifying poly(A) sites from ST-MOB, the expression level of each poly(A) site can be quantified by counting effective reads of the respective poly(A) site region (i.e., peak). Here we analyzed 3′ UTR APA genes. For genes with multiple poly(A) sites in 3′ UTR, RUD score (between 0 and 1) is used to measure the global trend of 3′ UTR length change of a gene. An RUD matrix was generated for ST-MOB data, with rows representing genes and columns representing spots. To mitigate the impact of high dropout rate, we proposed a KNN-based imputation method implemented in our stAPAminer package for recovering the RUD matrix (Materials and methods). First, we examined the clustering performance based on the comprehensive index for different *k* values from 2 to 100, and determined that the optimal *k* value for the KNN model is 10 (**Figure 3A**). After imputation with the KNN model (*k* = 10), we examined whether stAPAminer can efficiently recover the profile of APA usages. We applied UMAP visualization on the raw and the imputed RUD matrix to show differences between MOB layers. The 2D embeddings show that different layers became more distinguishable after imputation (Figure 3B). The inferred layer labels of spots based on the imputed RUD matrix are more consistent with the reference labels than those inferred from the raw RUD matrix (e.g., adjusted rand index (ARI): imputed = 0.643; raw = 0.547) (Figure 3C). Similar results were obtained using other three metrics including Jaccard, normalized mutual information (NMI), and Purity. Additional four metrics without relying on the reference labels, including Davies and Bouldin index (DBI) [38], Calinski-Harabasz (CH) [39], silhouette coefficient (SC) [40], and Dunn index [41], were used to quantitatively assess the spot separation. According to all these metrics, the imputed RUD matrix improves spot separation by recovering true signals of APA usages (Figure 3C). We further compared the spot-spot correlation in the same layer of the tissue section using the imputed and the raw RUD matrix. The median Pearson’s correlation of spot pairs in the same layer is only 0.311 to 0.501 using the raw RUD matrix, while stAPAminer greatly increased the spot-spot correlation of all the five layers (from 0.525 to 0.622) (Figure 3D). Previously, scDaPars [26] was proposed for identifying and quantifying APA events from scRNA-seq data. It first identifies APA events from scRNA-seq using DaPars [8], a tool for identifying APA events from bulk RNA-seq, and then utilizes a regression model to impute missing values in the APA usage matrix obtained from scRNA-seq. Although the underlying strategies are different, both stAPAminer and scDaPars can impute sparse APA signals. Here we used scDaPars to recover raw APA signals obtained from ST-MOB, and then compared the performance of scDaPars and stAPAminer in imputing APA signals. Regardless of the performance indicators used, stAPAminer outperforms scDaPars (Figures 3E and 3F), demonstrating again the superiority of stAPAminer in imputing highly sparse APA signals.

**Figure 3.**
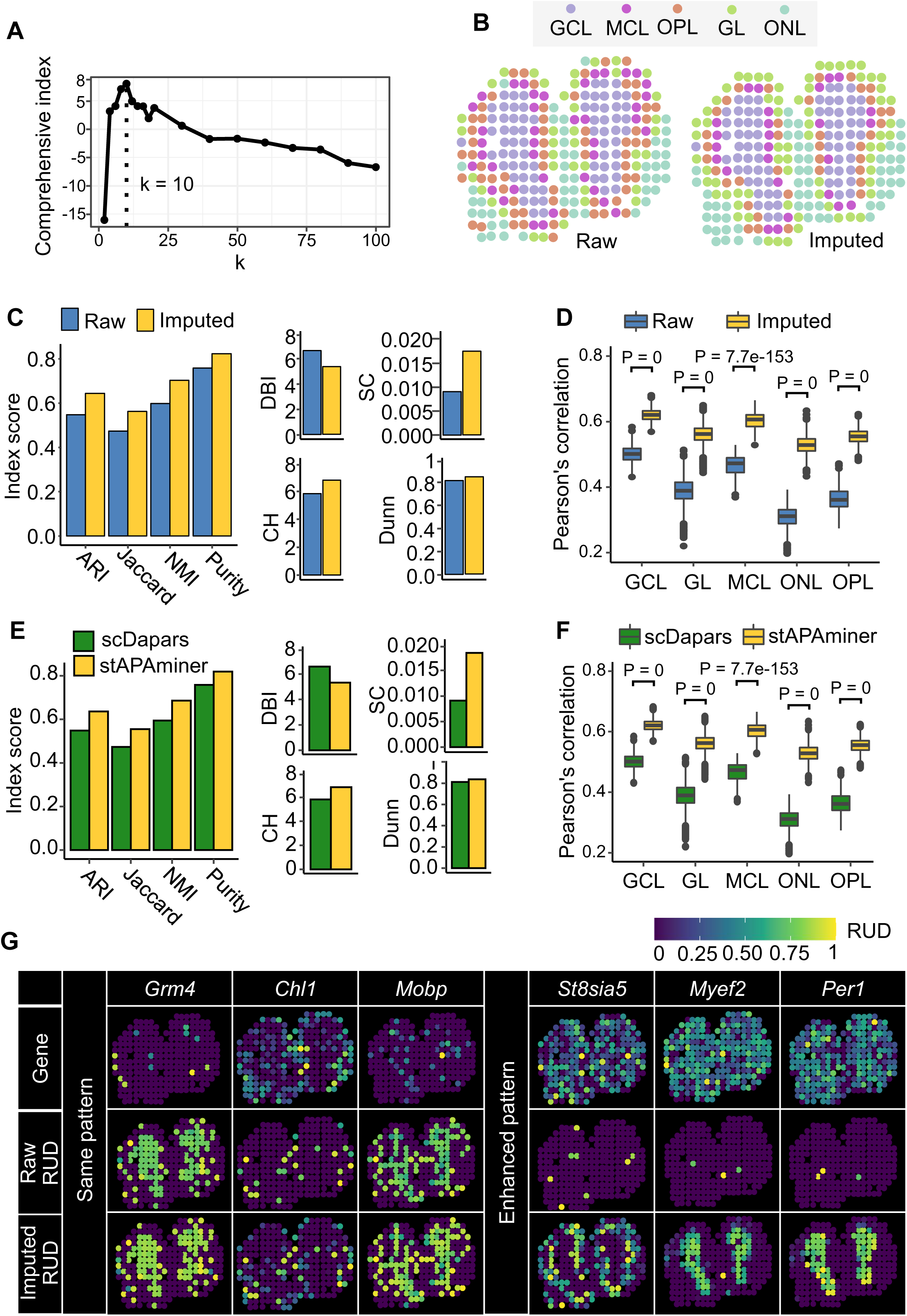
Validation of stAPAminer in imputing APA signals. **A.** The comprehensive index score with the increase of the *k* value. The vertical line marks the optimal *k* value (10), corresponding to the maximum comprehensive index score. B. Visualization of ST spots on the tissue image before (raw) and after (imputed) APA imputation. **C.** Evaluation of the performance of the imputation model. Four metrics were used for evaluating the performance in the context of clustering, including ARI, Jaccard, NMI, and Purity, and four internal validation metrics without relying on the reference labels were also used, including DBI, CH, SC, and Dunn. **D.** Boxplot showing Pearson’s correlations between spot pairs in each layer estimated using imputed and the raw RUD scores. For each layer, correlations of all pairwise spots were calculated. **E.** As in **C** except that the comparison was conducted between stAPAminer and scDaPars. **F.** As in **D** except that the comparison was conducted between stAPAminer and scDaPars. **G.** Spatial patterns of APA usages for representative genes using raw (middle) and imputed (bottom) RUD matrices. The top row shows the tissue image based on the scaled gene expression level of the respective gene. Color represents RUD scores or scaled gene expression levels (yellow, high; blue, low). ST, spatial transcriptomics; RUD, relative usage of distal poly(A) site; ARI, adjusted rand index; NMI, normalized mutual information; DBI, Davies and Bouldin index; CH, Calinski-Harabasz; SC, silhouette coefficient.

Next, we examined whether stAPAminer could reveal spatial APA usage patterns. We adopted SPARK [33] to identify genes with spatially variable usages of APA (i.e., SVAPA) (Materials and methods). For genes with strong spatial pattern of APA usage (e.g., the well-known layer-specific marker genes *Grm4* [42], *Chl1* [43], and *Mobp* [44]), the pattern from imputed data was concordant with that from the raw data (Figure 3G). Interestingly, the patterns of these genes are even more distinguishable according to the APA profile (either RUD or imputed RUD) than the raw gene-spot expression profile. Particularly, we also found cases where the raw RUD matrix exhibited a very weak signal that was enhanced in the imputed data (e.g., *St8sia5*, *Myef2*, and *Per1*) (Figure 3G). For example, *St8sia5* (ST8 alpha-N-acetyl-neuraminidase alpha-2, 8-sialyltransferase 5) is highly expressed on all layers, while no pattern was found according to either the gene expression profile or the raw RUD profile. In contrast, after imputation, distinct pattern was observed for this gene with much higher RUD scores (i.e., longer 3′ UTR) on mitral cell layer (MCL) or outer plexiform layer (OPL) than on other layers. Two 3′ UTR poly(A) sites were identified from ST-MOB for *St8sia5* (Table S1), both of which were supported by annotated poly(A) sites from PolyA_DB 3 [45]. *St8sia5* has been reported to induce the expression of ganglioside *GQ1b* and enhance neuronal differentiation via the *MAP* kinase pathway [46]. Another gene *Myef2* (myelin expression factor 2) is a transcriptional repressor of the myelin basic protein gene, which has been found usually upregulated in nerve sheath myxomas and schwannomas [47]. This gene also has two 3′ UTR poly(A) sites identified from ST-MOB, both of which are annotated in PolyA_DB 3. *Myef2* has a unique pattern according to the imputed RUD profile, with longer 3′ UTR on granular cell layer (GCL) than on other layers. Similar case was observed for *Per1* (period circadian regulator 1), which contributes to phasing molecular and electrical circadian rhythms in suprachiasmatic nucleus neurons to increase the robustness of cellular timekeeping [48]. These results show that the imputation strategy in stAPAminer can effectively mitigate the noise caused by the dropout phenomenon and help detect genes with distinct patterns of APA usage from the highly sparse and noisy ST data.

### stAPAminer reveals spatial dynamics of 3′ UTR APA usages from ST-MOB

Based on the imputed gene-spot RUD matrix, next we explored APA usage profiles in spatially defined domains within the olfactory bulb. The RUD profile well separates different morphological layers (**Figure 4A**). We detected 905 genes (403 non-redundant genes) with differential APA usage (i.e., DEAPA genes) between each pair of morphological layers (Table S2). For comparison, we also identified 1146 DE genes (DEGs) using only the gene-spot expression matrix (Table S3). Although the number of DEAPA genes and DEGs is comparable, the overlap between the two gene sets is very limited and a considerable number of genes are exclusively present in the DEAPA gene list. For example, between the GCL and MCL layer, only one common gene was found in the 78 DEGs and 54 DEAPA genes (Figure 4B). Similar case was observed between the GCL and olfactory nerve layer (ONL). These DEAPA genes were not recognizable by conventional gene expression analysis, while they represent genes with differential APA usage between layers. Particularly, among the 905 DEAPA genes, 111 genes were not present in the conventional gene-spot expression matrix. This may be due to that the scAPAtrap we used is highly sensitive in capturing poly(A) sites, even for lowly expressed genes, while these genes may failed to be detected in conventional gene expression analysis pipelines. Figure 4C shows two representative example genes, which showed clear spatial APA usage patterns but no gene expression patterns. *Adarb2* (also known as *ADAR3*) was extremely lowly and loosely expressed on GCL or ONL according to the gene expression profile, and no pattern was observed between these two layers. In contrast, it has two 3′ UTR poly(A) sites exclusively identified from ST-MOB (Table S1), showing differential APA usage between GCL and ONL. *Adarb2* encodes a catalytically inactive protein, mainly expressed in brain, thalamus, and amygdala, and may be associated with disorders such as amyotrophic lateral sclerosis [49]. *MAPK8IP3* (mitogen-activated protein kinase 8 interacting protein 3) encodes a protein that works like a motor in brain neurons, moving items along the axon of brain neurons [50]. *MAPK8IP3* is moderately expressed on both OPL and ONL without differential gene expression. Two 3′ UTR poly(A) sites were identified for this gene (Table S1), and differential APA usage was observed between OPL and ONL. Therefore, the APA profile potentially encodes complementary information that is absent or insignificant in conventional gene-spot expression profile, which contributes to distinguishing morphological layers. Next, we attempted to examine the presence of these DEAPA genes in known biologically important genes in the olfactory system. We complied representative genes in the olfactory system presented in previous studies or public resources (Table S4), including marker genes highlighted in the original study [30], cell type-specific marker genes provided in a recent scRNA-seq study on MOB [36], and genes related to the olfactory system from the Harmonizome database [51]. A moderate number of DEAPA genes (71 out of the 403 redundant genes) were present in the representative gene list, suggesting potential biological function of these APA genes. Indeed, the overlap is relatively limited, however, this is not unexpected as these DEAPA genes were identified based on spatial APA profiles independent of gene expression, while the compiled representative gene list is based on gene expression profile.

**Figure 4.**
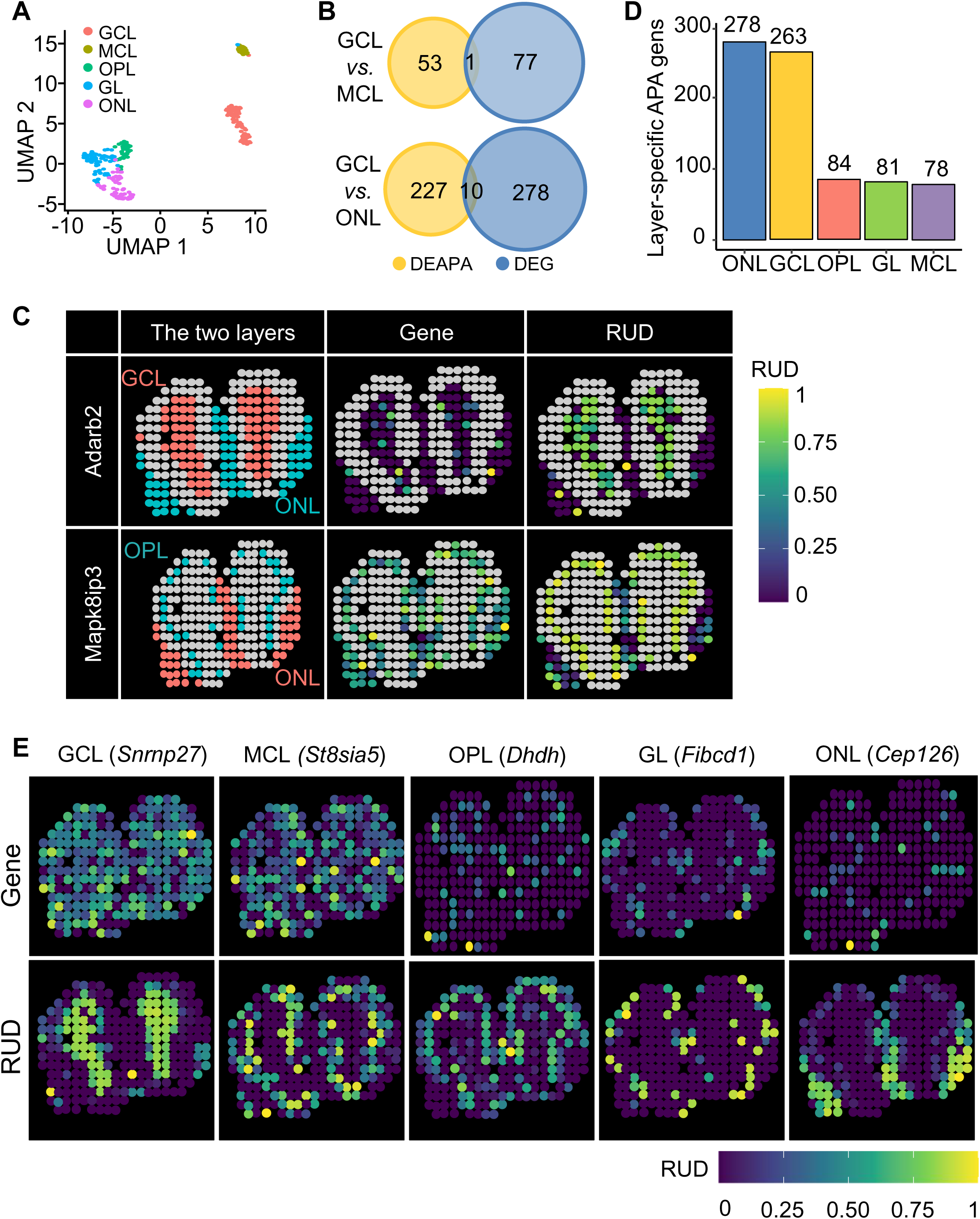
Application of stAPAminer to ST-MOB data for identification and analysis of genes with spatial APA pattern. **A.** UMAP plot showing the 2D-embeddings of spots on the five layers. **B.** Venn diagrams showing the overlap between DEAPA genes and DE genes by comparing GCL and MCL (top) and GCL and ONL (bottom). **C.** Two representative genes showing clear spatial APA usage patterns but no gene expression patterns. *Adarb2* is a DEAPA gene between GCL and ONL; *Mapk8ip3* is a DEAPA gene between OPL and ONL. Gene expression levels and RUD scores of spots on the respective layers are shown, while spots on other layers are colored as grey. **D.** Number of layer-specific APA genes on each layer. **E**. Representative layer-specific APA genes on the five layers, which showed clear spatial APA usage patterns but no gene expression patterns.

In addition to the DEAPA analysis, we further detected 784 layer-specific DEAPA (i.e., LSAPA) genes by comparing the RUD profile of a layer to all other layers (Table S5). Majority of these LSAPA genes (69%) were detected on layer GCL or ONL (Figure 4D). Figure 4E shows some representative LSAPA genes, which showed clear spatial APA usage patterns but no gene expression patterns. For example, *Snrnp27* and *St8sia5* are constitutively expressed across all layers, while they show distinct spatial APA usage pattern on GCL and MCL, respectively. *Snrnp27* has two poly(A) sites identified from ST-MOB (Table S1), and a previous study based on microarray found altered expression of this gene during progression of Alzheimer’s disease [52]. Additional, *Fibcd1*, a known chitin-binding receptor of the innate immune system, shows glomerular layer (GL) specificity according to both gene expression profile and APA profile, however, the layer-specific pattern obtained from the APA profile was more distinguishable. *Fibcd1* has recently been recognized as an evolutionarily conserved component of brain extracellular matrix, which is associated with a complex neurodevelopmental disorder [53]. Interestingly, another two genes, *Dhdh* and *Cep126*, are not detected from the gene-spot expression profile, however, they show layer-specific APA usage on OPL and ONL, respectively. *Cep126* is a regulator of microtubule organization at the centrosome, which has been found associated with diseases like include amyotrophy and monomelic [54]. It is a bit odd here that a gene is not expressed in any spot but its poly(A) sites can be detected. This is probably due to the different tools (or strategies) used for quantifying genes and poly(A) sites. Here the gene expression profile was obtained by conventional gene expression analysis pipeline, while the poly(A) site expression profile was obtained by scAPAtrap. scAPAtrap is very sensitive and can detect poly(A) sites for lowly expressed genes [25], while such genes may be undetectable by tools like Cell Ranger. Since the sum of expression levels of poly(A) sites in a gene can also be considered as the gene expression, we also summarized the read counts of all poly(A) sites in these two genes to represent their gene expressions, while no spatial pattern was observed for these two genes (Figure S3). These results indicate that the profile of APA usages encodes an additional layer of spatial information with higher resolution that is invisible in the gene expression profile.

Next, we adopted SPARK to identify SVAPA genes. Using the imputed RUD matrix as the input, a total of 133 genes with adjusted *P*-value < 0.05 were considered as SVAPA genes (Table S6). Similarly, we identified 772 SV genes with SPARK using the gene-spot expression matrix as the input (Table S7). Very limited overlap was found between these SVAPA genes and SV genes; only 16 genes are detected as both SVAPA and SV genes (Figure S4). This is not surprising in that SVAPA genes and SV genes were detected based on APA profile and gene expression profile, respectively. These wo independent groups of genes possess spatial patterns at the gene level and at the APA isoform level, respectively.

By combining DEAPA, LSAPA, and SVAPA genes, we obtained a comprehensive list of 654 non-redundant genes with spatial APA usage pattern (**Figure 5A**). A limited number of genes (101) were common in all the three gene sets, indicating that these three gene sets are complementary in reflecting the full landscape of spatial APA dynamics. We performed clustering on the combined gene set and obtained five major spatial expression patterns (Figure 5B): pattern I representing ONL; pattern II representing the combination of GL and MCL; pattern III representing the combination of GL, ONL, and OPL; pattern Ⅳ representing GCL; and pattern Ⅴ representing layers other than ONL. All the five patterns were clearly visualized via five representative genes, *Thg1l*, *Zfp983*, *Zfp974*, *Srnp27*, and *Srcap* (Figure 5C). For example, *Thg1l*, a tRNA-histidine guanylyltransferase 1-like protein associated with an autosomal recessive ataxia with abnormal neurodevelopment [55], shows generally higher RUD scores (e.g., longer 3′ UTR) on ONL. In contrast, *Srcap*, a *Snf2* related *CREBBP* activator protein associated with diseases like musculoskeletal defects and behavioral abnormalities, displays higher RUD scores on multiple layers except for ONL. Next, we performed functional enrichment analyses of the combined gene list with 654 genes. A total of 66 gene ontology (GO) terms were enriched in these genes at a false discovery rate (FDR) of 5% (Table S8). Many enriched GO terms are directly associated with functions related to olfactory bulb development, such as synapse organization, neuron differentiation, and nervous system development. These identified APA genes and GO terms highlight the benefits of spatial APA analysis with scAPAminer.

**Figure 5.**
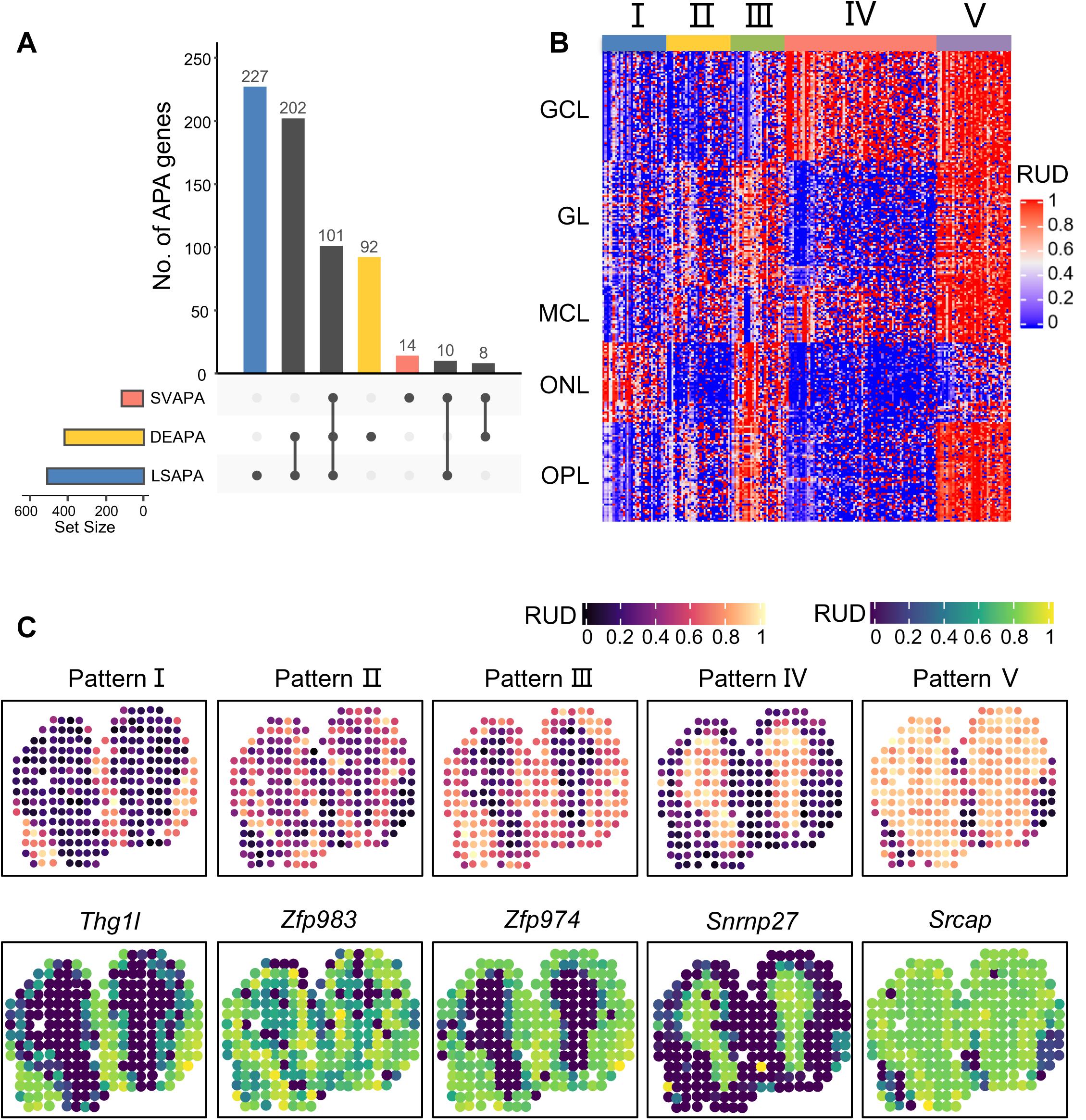
Combined analysis of genes with spatial APA usage pattern. **A.** Upset plot showing overlap of genes with differential APA usages (DEAPA), layer-specific APA genes (LSAPA) and spatially variable APA genes (SVAPA). **B.** Five major spatial APA patterns by clustering on the combined gene set. **C.** Representative tissue images of the five patterns (top) and the corresponding example genes for individual patterns (bottom). The tissue image for each pattern was generate by averaging RUD scores on each spot for all genes with the respective pattern.

### Spatial 3′ UTR APA patterns explored from additional replicates of ST-MOB demonstrate the robustness of stAPAminer

Next, we explored spatial APA patterns from two additional replicates of ST-MOB to evaluate the robustness of stAPAminer. We extracted poly(A) sites from two adjacent tissue sections (Replicate 5 and 12) of ST-MOB for the analysis of spatial APA dynamics. The profile of poly(A) site expressions of ST-MOB (which is Replicate 11) is highly similar to both replicates (Pearson’s correlation = 0.87 and 0.86) (Figure S5). Moreover, when repeating the imputation procedure on Replicate 5 (Figure S6) and Replicate 12 (Figure S7), we observed that our results were highly reproducible. For both replicates, different layers became more distinguishable after imputation (Figures S6A and S7A), and the imputation can improve spot separation or spot clustering (Figures S6B and S7B). These results demonstrate the robustness of the imputation method in stAPAminer. We also compiled DEAPA, SVAPA, and LSAPA genes for both replicates, resulting in 695 non-redundant genes for Replicate 5 (Table S9) and 556 for Replicate 12 (Table S10), respectively. For all the three replicates, much more LSAPA or DEAPA genes were identified than SVAPA genes (Figures 5A and S8). This may be because that different strategies were used for characterizing spatial APA dynamics — LSAPA or DEAPA genes were identified by comparing between layers while SVAPA genes were identified by inferring the spatial trend globally.

Although numbers of genes with spatial APA pattern were comparable among the three replicates, a limited number of consensus genes (364) were identified from two or more replicates, and a considerable number of genes were exclusively identified from one replicate (**Figure 6A**). The main reason may be attributed to the inherent nature of high sparsity and high noise of the ST data as well as the biological variance suffered from different replicates. Still we observed many interesting and reproducible cases (Figures 6B and S9). For example, differential APA usage between GCL and ONL was observed for *Adarb2* in Replicate 11 (Figure 4C), and a similar pattern was also observed in Replicate 12 (Figure 6B), while this gene was not detected in Replicate 5. Interestingly, this gene seems to also display a weak spatial gene expression pattern according to the tissue image, however, the pattern couldn’t be detected by spatial expression analysis (i.e., the gene was not present in Table S7). Similarly, *Mpp2*, a postsynaptic *MAGUK* scaffold protein [56], showed higher expression level or RUD score on GCL than other layers, while the pattern of APA usage is much more distinguishable than that of gene expression (Figure S9). *Mpp2* was also detected as SV gene by spatial expression analysis (Table S7). Particularly, the spatial pattern of APA usage was unobserved due to high dropout rate in Replicate 11, while the pattern was recovered after APA imputation and was highly consistent with that of Replicate 5. It is probable that some weak gene expression patterns may originate from noise or some genes may possess both patterns of spatial gene expression and APA usage, but with varied strength. We also found some genes whose gene expression was not detected from the gene-spot matrix while poly(A) sites and spatial APA pattern were detected from the imputed APA profile (e.g., *Dnal1* and *Tmem268*) (Figure S9). This reveals the high sensitivity of stAPAminer in identifying and imputing APA profile for genes which may be missed by conventional gene expression pipelines. Taken together, these results demonstrate the ability and robustness of stAPAminer in identifying spatial APA patterns.

**Figure 6.**
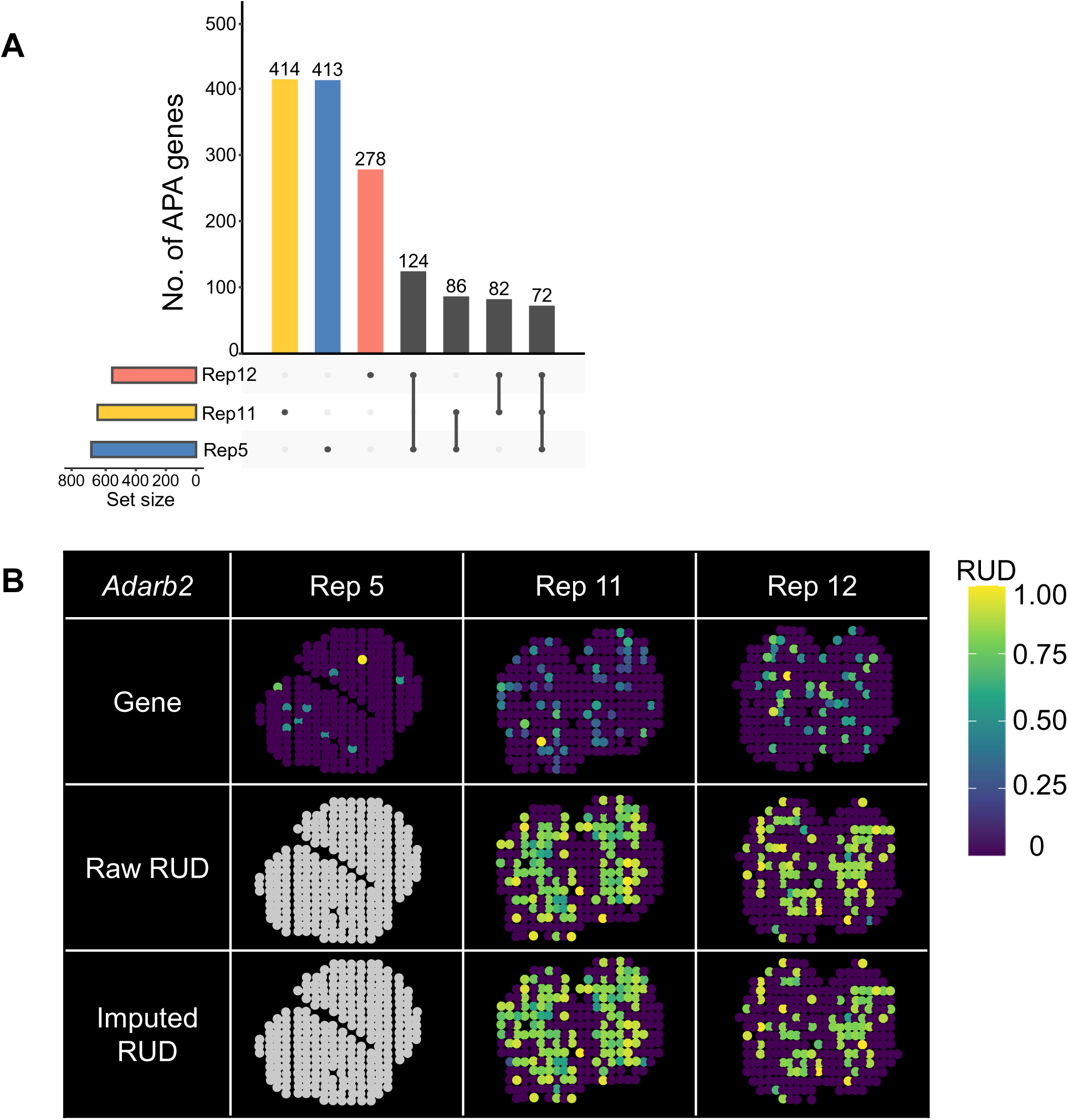
Analysis of spatial APA dynamics from additional two replicates of ST-MOB with stAPAminer. **A.** Upset plot showing overlap of genes with spatial APA pattern among the three replicates (Rep 5, Rep 11, and Rep 12). **B.** An example gene (*Adarb2*) with differential APA usage between GCL and ONL. Tissue images based on the scaled gene expression level (top), raw RUD scores (middle) and imputed (bottom) RUD matrices were shown. Color represents RUD scores or scaled gene expression levels (yellow, high; blue, low).

### stAPAminer reveals extensive spatial dynamics of intronic APA usages from ST-MOB

Having demonstrated spatial dynamics of 3′ UTR APA, we next sought to explore spatial patterns of intronic APA from ST-MOB. First, we obtained a matrix of intronic APA usage (Materials and methods), containing 4011 genes present in 260 spots. The matrix was then imputed with the KNN model in stAPAminer (*k* = 10). The layer clustering result based on the imputed intronic APA signals is similar to that based on 3′ UTR APA signal (ARI: intronic APA = 0.605, 3′ UTR APA = 0.643) (**Figures 7A** and 3B). Based on the imputed intronic APA signal matrix, we further identified 601 non-redundant DEAPA genes (Table S11), 386 LSAPA genes (Table S12), and 226 SVAPA genes (adjusted *P*-value < 0.05) (Table S13). Combining these three sets of genes, 669 non-redundant genes with spatial APA usage patterns were obtained, among which 130 genes also show spatial 3′ UTR APA patterns (Figure 7B). Next, we clustered these genes based on their intronic APA profiles and obtained five major spatial APA patterns (Figure 7C): pattern I representing ONL; pattern II representing the combination of GL and MCL; pattern III representing the combination of GL, ONL, and OPL; pattern Ⅳ representing GCL; and pattern Ⅴ representing layers other than ONL. These five patterns were clearly visualized via five representative genes, *Adgrg6*, *Mia3*, *P3h2*, *Sema3c*, and *Rapgef2* (Figure 7D). For example, *Adgrg6* (adhesion G protein-coupled receptor G6) is a protein-coding gene that is essential for normal differentiation of promyelinating Schwann cells and normal myelination of axons [57]. *Mia3* (MIA SH3 domain ER export factor 3) is involved in cell migration related to sprouting angiogenesis [58]. *P3h2* (prolyl 3-hydroxylase) catalyzes the post-translational formation of 3-hydroxyproline on collagens [59]. *Sema3c* (semaphorin 3C) acts as an attractant for growing axons and thus plays an important role in axonal outgrowth and axon guidance [60]. *Rapgef2* (rap guanine nucleotide exchange factor 2) is involved in cAMP-induced Ras and Erk1/2 signaling, leading to sustained inhibition of long-term melanogenesis by reducing dendritic elongation and melanin synthesis [61]. Functional enrichment analysis of the combined gene list revealed that these genes were associated with the olfactory system and most of them were directly related to the structure and regulation of synaptic organization (Table S14). In contrast, although genes with spatial 3′ UTR APA patterns were also associated with olfactory bulb development, they were enriched in synaptic organization, neuronal differentiation, and nervous system development (Table S8). These results indicated extensive spatial dynamics of intronic APA usages, while genes with spatial intronic APA patterns are distinct from those with spatial 3′ UTR APA patterns.

**Figure 7.**
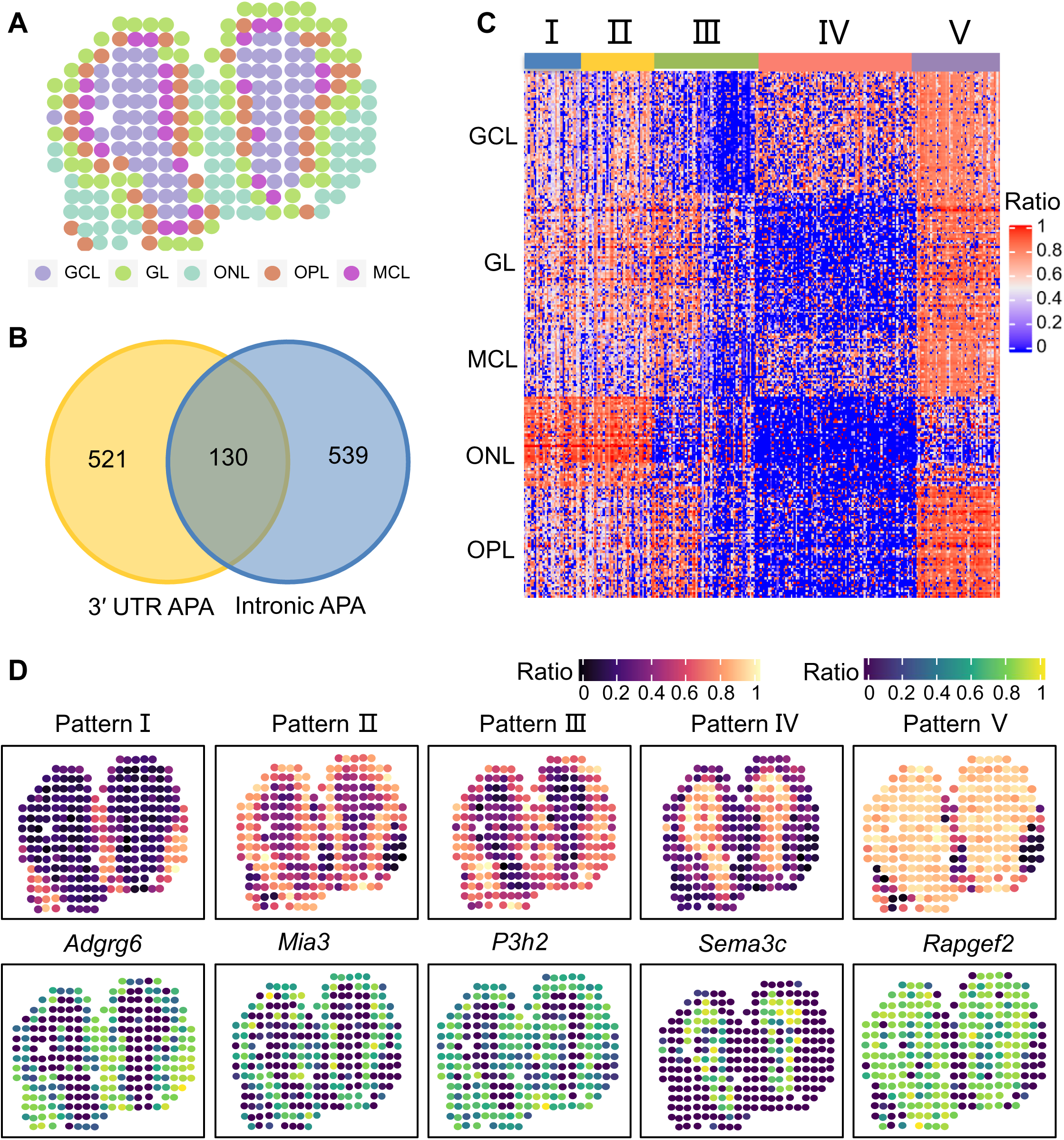
Analysis of spatial patterns of intronic APA from ST-MOB. **A.** Visualization of ST spots on the tissue image after intronic APA imputation. **B.** Venn diagram showing the overlap of 3′ UTR APA genes and intronic APA genes with spatially APA patterns. **C.** Five major spatial APA patterns by clustering on the combined gene set. **D.** Representative tissue images of the three patterns (top) and the corresponding example genes for individual patterns (bottom). The tissue image for each pattern was generate by averaging intronic ratio scores on each spot for all genes with the respective pattern.

### The KNN-based imputation model in stAPAminer is robust to data with different dropout rates and spot sizes

The KNN-based imputation method was embedded in our stAPAminer package for recovering the sparse RUD matrix; the most important parameter in the model is the *k* value. Here we evaluated the influence of spot size of the RUD matrix on the choice of the *k* value. First, we examined the influence of the number of dropout spots on the *k* value. We set the percentages of dropout spots of the MOB Replicate 11 from 10% to 90% by masking all genes in randomly selected spots while keeping the spot number among layers unchanged. Then the clustering performance based on the comprehensive index was evaluated under different *k* values at each dropout rate. In general, the larger the dropout rate is, the larger the optimal *k* value is (**Figure 8A**). This may be because when the dropout rate increases, more adjacent spots are needed to obtain sufficient information to achieve comparable clustering performance. However, even with an extremely high dropout rate (e.g., > 50%), a *k* value within 20 can basically yield moderately high performance. Next, to evaluate the influence of the spot size on the *k* value, we randomly sampled 20% to 90% spots from the total of 260 spots while keeping the proportion of spot number among layers unchanged, and then tested the performance of different *k* values. Regardless of different spot sizes, the optimal *k* value is around 10 (Figure 8B). After reaching the optimal *k* value, the performance decreases gradually with the increase of *k* value, which may be because that a larger *k* value may introduce greater biases by borrowing the information of spots from other layers. Generally, our model has good robustness to different *k* values. It is recommended to use a smaller *k* value with equally high comprehensive index score, which can ensure high performance and avoid introducing the biased information of spots from other layers due to a larger *k* value.

**Figure 8.**
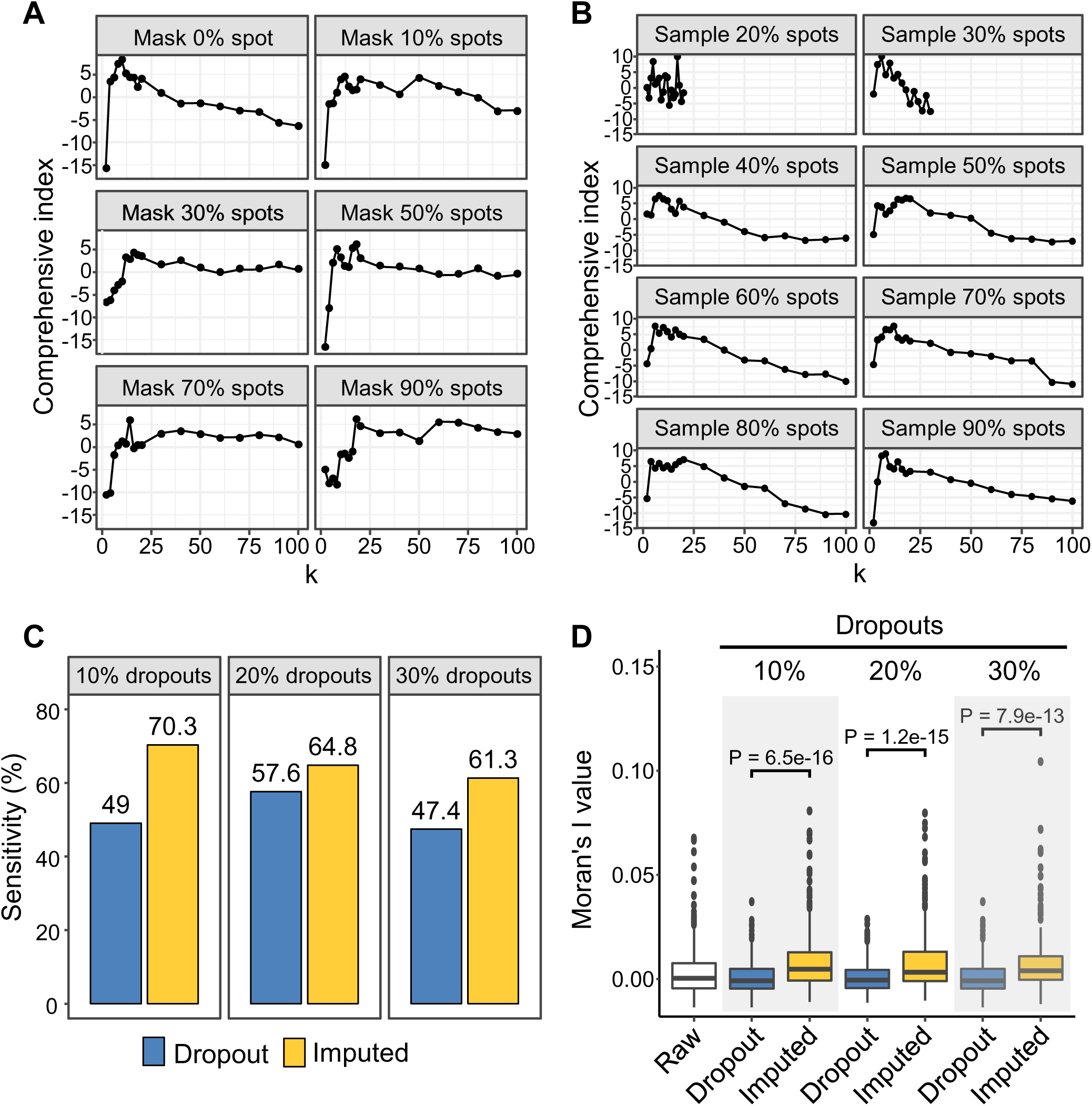
Evaluation of the KNN-based imputation model using RUD matrix with varied spot sizes and dropout rates. **A.** The value of the comprehensive index with the increase of k value under different number of dropout spots. The percentages of dropout spots of the were set from 10% to 90% by masking all genes in randomly selected spots while keeping the spot number among layers unchanged. **B.** The value of the comprehensive index with the increase of k value under different number of sampled spots. 20% to 90% spots were randomly sampled while keeping the proportion of spot number among layers unchanged. **C.** Sensitivity of the KNN-based imputation model under RUD matrices with different dropout rates. The dropout rate of the raw RUD matrix was increased by 10%–30% through randomly masking values in the matrix. Genes with spatial APA pattern present in at least two replicates of Replicate 5, 11, and 12 were used as the true reference. Spatial APA patterns from the RUD matrix before and after imputation were identified to obtain sensitivity, respectively. **D.** Moran’s I value of spatial APA patterns identified from RUD matrices with different dropout rates. The RUD matrix was processed as in **C**. For all plots in this figure, the RUD matrix of MOB Replicate 11 was used, which contains a total of 260 spots.

Next, we examined whether the KNN-model in stAPAminer is overfitted or not, and evaluated the sensitivity of stAPAminer in recovering spatial APA patterns. To the best of our knowledge, this is the first study on the analysis of spatial APA pattern and there is no gold standard about genes with spatial APA patterns, it is thus difficult to validate whether an identified spatial APA pattern is true or not. Alternatively, we try to compile a silver standard dataset containing genes with reproducible spatial APA pattern. Briefly, we first identified genes with spatial APA pattern in each of the three replicates (Replicate 5, 11, and 12) of raw ST-MOB data analyzed in this study, and established a list of 401 genes with spatial APA pattern present in at least two replicates. These genes were considered as authentic instances for further evaluation. Assuming that our KNN-based imputation model is overfitted, the accuracy of spatial APA patterns identified from the recovered RUD matrix would be lower than that of the raw RUD matrix. Accordingly, we mimic data with higher sparsity to verify whether the KNN-base imputation model is overfitted. We increased the dropout rate of the already sparse raw RUD matrix by 10%–30% through randomly masking values in the matrix. Then we used the KNN-base model to recover the extremely sparse RUD matrix. We identified spatial APA patterns from the RUD matrix before and after imputation respectively, and calculated sensitivity according to the compiled sliver standard. Regardless of the sparsity of the data, the sensitivity of pattern recognition on the imputed data is much higher than that before imputation (Figure 8C). It should be noted that the silver standard data we constructed only represents true instances, whereas spatial APA patterns not in the silver standard are not necessarily false. Since there is no gold standard for false instances, only true positives (TP) and false negatives (FN) rather than false positives (FP) or true negatives (TN) can be calculated. Therefore, here we only calculated sensitivity to evaluate the results. Moreover, we used Moran’s I [62] value to measure the significance of the identified spatial patterns, and found that the spatial autocorrelation of the patterns obtained after imputation is significantly improved (Figure 8D). These evaluation results argue very strongly even for highly sparse data, our KNN-based imputation model is unlikely overfitted and can effectively enhance or recover the RUD signal, thereby greatly improve the sensitivity of spatial pattern recognition.

## Discussion

ST technologies have the advantage of revealing an unbiased map of transcripts over complex tissues and cell cultures. Established spatially resolved approaches can detect genes with localized expression patterns at single-cell resolution. However, these approaches mainly focused on gene-level analysis rather than isoform-level analysis. Consequently, these studies can neither measure isoform usages in a spatially defined tissue region nor detect significant spatial isoform usage in a spatial context. There is a clear need for developing new methods for exploring isoform usages from spatially resolved transcriptional data. The rich data generated by ST experiments promise to isoform-level analysis, which complements conventional gene-level studies using ST data. We present an innovative method, stAPAminer, for analyzing spatial APA dynamics in spatially resolved transcriptomic studies, which allows anchoring the APA isoform usage in a spatial view and reveals important spatial expression patterns at the isoform resolution. The integration of APA isoform usages and gene expression in the spatial context brings the promise of establishing spatial maps of APA dynamics. Our work furthermore defines an atlas of APA events with distinct spatial APA usage patterns, which supplements spatially variable genes for capturing the spatial complexity of MOB subregions. We have verified the identified genes with significant spatial APA usage in various ways and demonstrated the improvement of layer classification by integrating the spatial APA information.

Limited by the sequencing depth, gene expression profile obtained from the ST data is usually in sparse forms, with ubiquitous low counts and zeros. Moreover, the obtained APA profile is sparser than the already sparse gene expression profile, which presents a huge computational challenges for studying the spatial patterns of APA isoform usage. Our stAPAminer pipeline incorporates a KNN-based imputation model for recovering APA signals, which mitigates the sparsity of the APA isoform matrix and yields stronger spatial patterns. The evidence for APA isoforms adding to cellular diversity has been bolstered by the APA signal imputation (Figure 3). After imputation, the signal of genes with spatial pattern of APA usage was retained or greatly enhanced (Figure 3F). Additional analyses using data with varied spot sizes and different number of dropout spots and genes also demonstrate that the KNN model is robust to different *k* values and can greatly improve the sensitivity of spatial pattern recognition (Figure 8). stAPAminer has several desirable features for mitigating high sparsity of the APA profile. First, scAPAtrap [25] used in our pipeline is capable of accurately identifying all potential poly(A) sites, even those with low read coverage, by incorporating the strategy of peak identification and poly(A) read anchoring. Second, the KNN-based imputation model in stAPAminer can efficiently impute missing values by clustering spots based on gene signatures rather than the APA profile alone. This promises the inference of associations between ST spots by borrowing information from the gene expression profile of the ST data, which avoids the lack of information or over-fitting when using the APA data alone. Third, multiple rounds of iterations are performed during the imputation process, which enables gradually inclusion of newly imputed information.

Using stAPAminer, we identified three kinds of genes with spatial APA dynamics, including genes with differential APA usage between each pair of morphological layers (DEAPA), layer-specific DEAPA (LSAPA) genes, and genes with spatially variable APA patterns (SVAPA). Moreover, we explored spatial patterns of both 3′ UTR APA and intronic APA. A total of 654 non-redundant 3′ UTR APA genes from the combined list of DEAPA, LSAPA and SVAPA genes were obtained from the ST-MOB data (Figure 5A), representing five major spatial expression patterns (Figure 5B). Similarly, 601 intronic APA genes with spatial pattern were obtained, however, interestingly, these genes are distinct from those with spatial 3′ UTR APA patterns (Figure 7). We provided several lines of evidence to validate these genes with spatial APA dynamics. First, the poly(A) sites identified from the ST-MOB data are of high confidence, which are supported by annotated poly(A) sites and possess typical poly(A) signals (Figure 2). Second, we examined the presence of these genes in the list of representative biologically important genes in the olfactory system from several previous studies [30, 36] (Table S4). Third, enriched GO terms of these spatially variable APA genes are directly associated with functions related to olfactory bulb development (Table S8). Fourth, when overlaying the APA signal of a spatially variable APA gene on the tissue images, well-defined spatial patterns were revealed (Figures 3–5). Fifth, spatial APA patterns of many genes are reproducible among replicates (Figures 6 and S9). It is worth noting that spatially variable APA genes identified by stAPAminer do not necessarily present in genes obtained from conventional gene expression studies, however, these lines of evidence provide significant clues to the functional importance of the identified APA genes.

In comparison with spatial patterns obtained by spatial expression analysis, we observed several interesting cases of spatial APA patterns. We found some highly expressed genes with distinguishable spatial APA patterns but without any spatial gene expression pattern (e.g., *St8sia5*, *Myef2*, and *Per1*) (Figure 3F). In addition, some genes present both detectable spatial APA patterns and gene expression patterns, with the APA pattern being more distinguishable (e.g., *Mpp2*) (Figure S9). There are also some genes seem to have spatial gene expression pattern but fail to be computationally detected by existing tools, while they have apparent spatial APA patterns (e.g., *Adarb2*) (Figure 6B). These different groups of genes suggest the presence of spatially variable patterns at the APA isoform level independent of the gene expression profile. In particular, some genes are not found in the gene-spot matrix while spatial APA patterns were detected (e.g., *Dnal1* and *Tmem268*) (Figure S9). This result is unlikely to be due to noise or random bias, as we observed the same pattern in at least two replicates. The main reason may be the different pipelines used to obtain the gene-spot expression matrix and the gene-spot RUD matrix. The tool scAPAtrap [25] used in this study has been proven to be highly sensitive and is able to identify poly(A) sites in extremely low-expressed genes, while these low-expressed genes may failed to be detected in conventional gene expression analysis pipelines. Moreover, the APA signals are amplified through the imputation process of stAPAminer, which further contributes to the successful identification of spatial APA patterns. Taken together, these results demonstrate the effectiveness and robustness of stAPAminer in identifying different spatial APA patterns. It would also be necessary in the future to benchmark different methods on obtaining gene expression matrices and/or RUD matrices, as well as to propose methods that can mitigate discrepancies among replicates for identifying more reproducible spatial patterns.

In summary, we present a detailed view of spatial APA usage across MOB regions with our proposed toolkit stAPAminer. Our stAPAminer approach is, to our knowledge, one of the first computational approach for exploring transcriptional atlas of spatial APA patterns with spatial resolution. stAPAminer employs the power of ST to explore genome-wide spatial patterns of APA usage variation at the isoform resolution, establishing an additional layer of gene expression. The combination of both spatial maps of gene expression and APA usage will allow us to deduce a more comprehensive set of genes summarizing the spatial and cellular information, redefining the overall transcriptome complexity of a tissue.

## Materials and methods

### Data

The input of stAPAminer is a poly(A) site matrix with each row being a poly(A) site and each column being a spot. Currently, there are several ST strategies such as MERFISH [28], seqFISH [29], and ST through spatial barcoding [30]; among them, APA sites can only be identified from the spatial barcoded ST data for now, using existing scRNA-seq tools like scAPAtrap or Sierra. However, if APA sites can be identified from other kinds of ST data in the future, stAPAminer would also be applicable to those ST data. In this study, the spatial barcoded ST data of MOB [30] was used. We downloaded gene-expression measurements of the MOB data from Spatial Transcriptomics Research (http://www.spatialtranscriptomicsresearch.org), which were collected on spatial locations known as spots. The corresponding array oligonucleotides with positional barcodes were also obtained. Following the studies of SpatialDE [31] or SPARK [33], the ‘MOB Replicate 11’ file was mainly used, which consists of 16,218 genes measured on 262 spots. We further retained spots with ten or more read counts, resulting in 260 spots. Two additional replicates of MOB (Replicate 5 and 12) were also used for validation. The raw sequencing data (accession number: SRR3382373) was downloaded from NCBI SRA. Lengths of barcodes and unique molecular identifiers (UMIs) are 18 nt and 9 nt, respectively. The genome assembly (version GRCm3) and the latest genome annotation of mouse were downloaded from Ensembl.

### Identification and quantification of APA sites from ST data

The process of extracting APA sites from ST data is similar to that of scRNA-seq data. The raw ST data are double stranded, including read 1 and read 2. First, barcodes and UMIs were extracted from read 1 and then umi_tools [63] was adopted to append the barcode and UMI information to the sequence header of read 2 to generate a new read 2 FASTQ file. Reads from the read 2 file were aligned to the reference genome using STAR [64] and only uniquely mapped reads were retained. After mapping, PCR duplicates were removed by the *dedup* function in umi_tools and one read per UMI for each genomic coordinate was remained. Then scAPAtrap [25] was used to identify poly(A) sites from these mapped reads. Poly(A) sites were annotated with information, such as genomic region and gene, with the movAPA package [65] using the R annotation package ‘txdb.musculus.ucsc.mm10.knowngene’. Similar to previous studies [66–69], 3′ UTRs of annotated genes were extended by 1000 bp to recruit intergenic poly(A) sites which may be originated from authentic 3′ UTRs.

### Quantification and imputation of spatial APA usages

We calculated the relative usage of the distal poly(A) site (RUD) for each 3′ UTR APA gene from the ST data using the *movAPAindex* function in movAPA [65]. Briefly, only 3′ UTR APA genes that contain at least two poly(A) sites in 3′ UTR were retained. The distal poly(A) site is the one which is farthest from the start codon among all 3′ UTR sites. The RUD score of each gene is the ratio of the expression level of the distal poly(A) site to the sum of the expression levels of all poly(A) sites located in 3′ UTR. The RUD value is between 0 and 1. A larger RUD value of a gene in a spot indicates the higher usage of the distal poly(A) site of this gene in this spot (i.e., 3′ UTR lengthening). If there is no poly(A) site expressed in a gene for a given spot, then the respective RUD value is called a dropout. Finally, an RUD matrix is obtained for a ST dataset, with each row denoting a gene and each column denoting a spot.

To explore the spatial patterns of intronic APA, we filtered poly(A) sites that are supported by at least three reads and present in at least three spots, and retained genes with multiple poly(A) sites and at least one intronic poly(A) site. We calculated the ratio for each intronic poly(A) site as the expression level of the respective site to the sum of expression levels of all sites in the same gene. The highest ratio of intronic poly(A) site(s) was considered as the intronic APA usage of a gene. The ratio value is also between 0 and 1. A larger ratio of a gene in a spot, indicates the higher usage of the intronic poly(A) site of the gene in this spot.

Due to inherent technical issues of spatial RNA-seq and lower expression at the APA isoform level than at the gene level, the APA usage matrix (RUD or ratio) is even sparser than the gene-spot expression matrix. Although the APA usage matrix contains usage information at the APA isoform level and has a higher resolution than the gene expression matrix, it was obtained from only APA genes, consequently, it does not store as many genes as the gene-spot expression matrix. To mitigate the high sparsity of the APA usage matrix, we designed a KNN-based imputation model and introduced the gene-spot expression matrix to obtain the correlation between spots to better impute the APA usage matrix. Given a gene-spot matrix *G* with *n* genes and *m* spots, first the matrix is scaled (including data centering and standardization). Next, the Euclidean distance between each two spots is calculated to obtain a spot-spot distance matrix *D* (Eq. 1). Then a ranking matrix 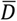 can be obtained, which is a spot-spot matrix storing the spot indexes for each spot in descending order according to the distance between all other spots and the respective spot. 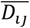 representing the spot index of the *j*^th^ closest spot for the *i*^th^ spot.

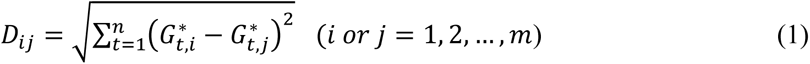

Here, 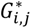 is the scaled gene expression level of gene *i* in spot *j*.

Given an APA usage matrix *R* with *h* genes and *m* spots, first values for those genes not expressed in the gene-spot matrix *G* are set as 0. Next, for each spot, nearest *k* spots (default *k* = 10) are selected according to the ranking matrix 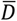. The nearest *k* spots for any spot *i* (i.e., the *i*^th^ row in 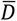) are the first *k* columns of the *i*^th^ row in the ranking matrix. And then for a gene with missing APA usage in matrix *R*, the average APA usage of this gene in these *l* spots is calculated for imputation (Eq. 2). Of note, only spots with a non-zero APA usage for the gene were counted.

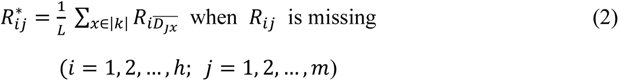

Here, 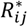 is the imputed APA usage for gene *i* in spot *j*; |*k*| denotes the set of nearest *k* spots with an APA usage for the respective gene; *L* is the number of spots in |*k*|.

After the first round of imputation, a small part of APA usage values remain missing when the respective genes are not expressed in any of the *k* nearest spots. We then repeated the above imputation step (Eq. 2) and performed multiple iterations until no missing value was filled. In each iteration, the average value of the APA usage of neighboring spots is calculated according to the newly imputed APA usage matrix. After the iteration process, there is still a small number of APA usage are missing, we set them to 0 for subsequent analysis.

We adopted several performance indicators to evaluate our KNN-based imputation model. First, we manually obtained the true layer where each spot is located by combining the spatial information of each layer with the Hematoxylin & eosin stained brightfield image of MOB slices. These true labels of layers were used as reference. To examine whether the imputed APA usage matrix better reflects the true relationship between spots than the raw APA usage matrix, first we adopted the *FindClusters* function (resolution = 5) in Seurat [70] to cluster spots based on the APA usage matrix and used four metrics to evaluate the performance in the context of clustering, including the ARI, Jaccard, Purity, and NMI. The ARI score ranges from −1 to 1, and scores of Jaccard, Purity, and NMI range from 0 to 1, with the higher value reflecting the better performance. In addition, four internal validation metrics, including DBI [38], CH [39], SC [40], and Dunn index [41], were employed to quantitatively assess the consistency of a clustering structure, which are independent of clustering methods or priori knowledge about true labels. A smaller DBI score or higher CH, SC, or Dunn score indicates better separation among clusters. Moreover, we calculated the Pearson correlation coefficient between each two spots under the same layer to examine whether the correlation of spots on the same layer after imputation is higher than that before imputation.

The most important parameter in our KNN-based imputation model is the *k* value. We proposed a strategy to determine the optimal *k* value based on a comprehensive index. Briefly, given a set of *k* values, the APA usage matrix is imputed under each *k*. Then clustering is performed on the imputed matrix to obtain eight clustering index values, including the four internal validation metrics (DBI, SC, CH, and Dunn) and the four external metrics (ARI, Jaccard, NMI, and Purity). Next, the Z-score is calculated for each index and the sum of all Z-score values is considered as the comprehensive index. Finally, the *k* value with the highest comprehensive index score is regarded as the optimal *k*.

### Identification of genes with spatial patterns of APA usage

To fully explore spatial patterns of APA usage, we identified three kinds of APA genes, including DEAPA, LSAPA, and SVAPA genes. DEAPA genes are genes that exhibit differential APA usage between two layers, which are analogue to DEGs in conventional gene expression studies. LSAPA genes are genes that exhibit layer-specific APA usage. SVAPA genes are genes that exhibit distinct spatial APA usage patterns in the global spatial context, which are analogue to SV genes in conventional ST studies. We followed the Seurat tutorial [70] to cluster the spots based on the APA usage matrix and obtained five clusters of spots (equal to the number of layers). These clusters were annotated as GCL, MCL, OPL, GL, and ONL according to the hematoxylin-and-eosin staining image. To identify DEAPA genes, we used the *FindMarkers* function in Seurat with the APA usage matrix as input. This function identifies DEAPA genes between two groups of spots using a Wilcoxon Rank Sum test. Genes with adjusted *P*-value < 0.05 and log_2_(FC) > 0.5 were considered as DEAPA genes. We also identified LSAPA genes with *FindMarkers* by comparing spots in a layer to all spots in remaining layers.

In addition, we also attempted to detect genes with spatial patterns of APA usage variation, i.e., SVAPA genes. Different from DEAPA genes that show differential APA usages between two groups of spots, SVAPA genes mark distinct spatial APA usage patterns in the global spatial context. Currently there is no tool available for identifying global spatial patterns from ratio type data, but there are several tools for identifying SV genes, which are hopeful to be applied to the APA usage matrix. Among them, SPARK [34] utilized a non-parametric method to effectively detect spatially expressed genes from large spatial transcriptomic data, which controls type I error and produces high power. SPARK was proven as ten times more powerful than other existing methods [34]. Because SPARK was designed for count data, we transformed the APA usage values which are between 0 and 1 into count data by taking 10 as the base (for 3′ UTR APA) or by log_2_ conversion (for intronic APA). The transformed APA usage matrix was then used as the input for SPARK and genes with adjusted *P*-value < 0.05 were considered as genes with spatial APA usage patterns. Moreover, we have implemented a unified interface in stAPAminer for importing SVAPA results, which facilitates users to incorporate results from other tools upon the availability of more dedicated tools in the future.

After obtaining DEAPA genes, LSAPA genes, and SVAPA genes, we combined these three sets of genes without redundancy to compile a unique gene set with dynamic APA usages in a spatial context. We then adopted k-means to cluster these genes into ten groups based on their APA usage profile, using the *kmeans* function of the R package stats with arguments ‘iter.max = 1e+9, nstart = 1000’. Next, the mean APA usage profile for each group of genes was calculated for each spot; each group was considered as one spatial APA pattern. We selected five groups with most distinguished spatial APA usage patterns. We take the selected spatial APA usage patterns as samples, calculate the Pearson’s correlation to measure the similarity between each gene in the gene set and the sample, and select the genes with Pearson’s correlation > 0.5 and *P*-value < 0.05 as the representative of the spatial pattern genes.

## Data availability

Our implementation of stAPAminer is available at the repository (https://github.com/BMILAB/stAPAminer).

## CRediT author statement

**Guoli Ji**: Investigation, Methodology, Data curation, Writing - review & editing. **Qi Tang**: Data curation, Software, Visualization, Writing - review & editing. **Sheng Zhu**: Data curation, Software, Visualization; **Junyi Zhu**: Data curation, Formal analysis. **Pengchao Ye**: Formal analysis. **Shuting Xia**: Formal analysis; **Xiaohui Wu**: Conceptualization, Writing - original draft, Writing - review & editing, Supervision, Project administration, Funding acquisition. All authors read and approved the final manuscript.

## Acknowledgments

This work was supported by the National Natural Science Foundation of China (Grant No. 61673323 to XW, 61573296 to GJ, and 81901287 to SX) and Suzhou City People’s Livelihood Science and Technology Project (Grant No. SYS2020086 to S.X.).

## Supplementary materials

**Figure S1.**
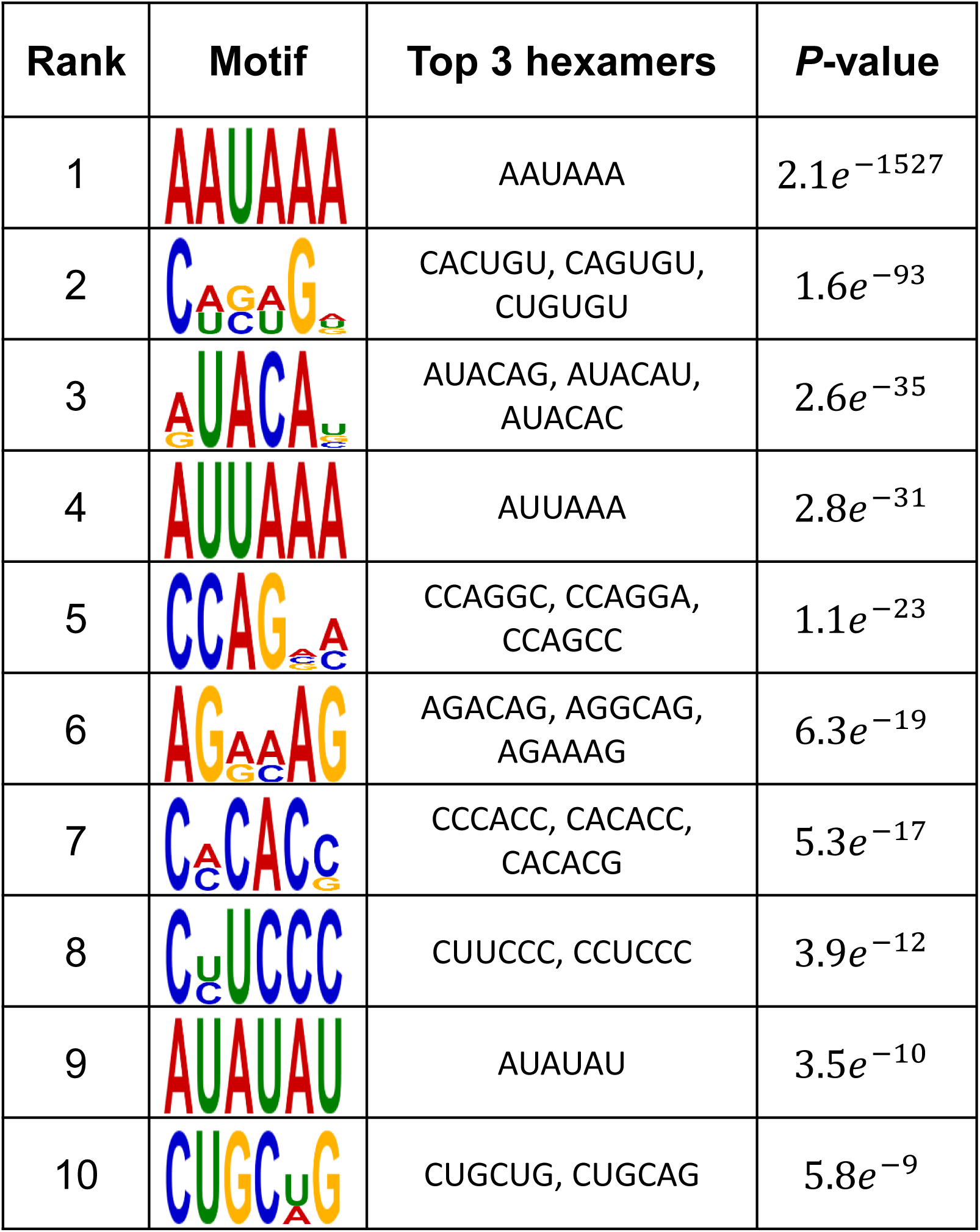
Motif identified in the upstream 50 nt region of the poly(A) site. Motifs were identified by MEME for 3′ UTR poly(A) sites. Top ten motifs, each with three most significant hexamers, were shown.

**Figure S2.**
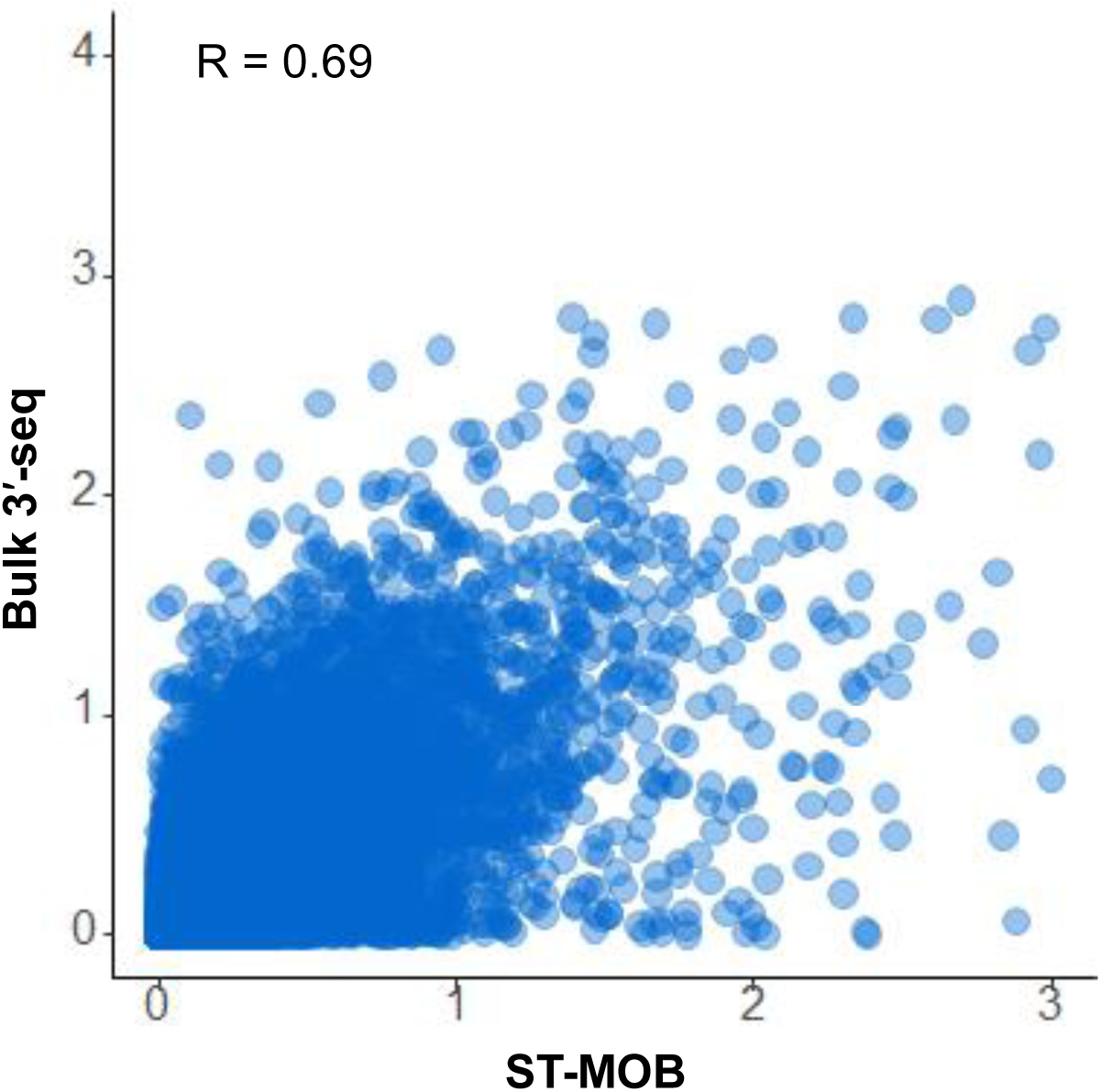
Scatter plot showing the correlation of poly(A) site expression profiles obtained from ST-MOB and bulk 3′-seq. Each dot is one poly(A) site and the axis is the natural-log scaled read counts. The Pearson’s correlation is indicated in the plot. Here the bulk 3′-seq data contains a total of 32 neural-related samples from the PolyASite 2.0 database.

**Figure S3.**
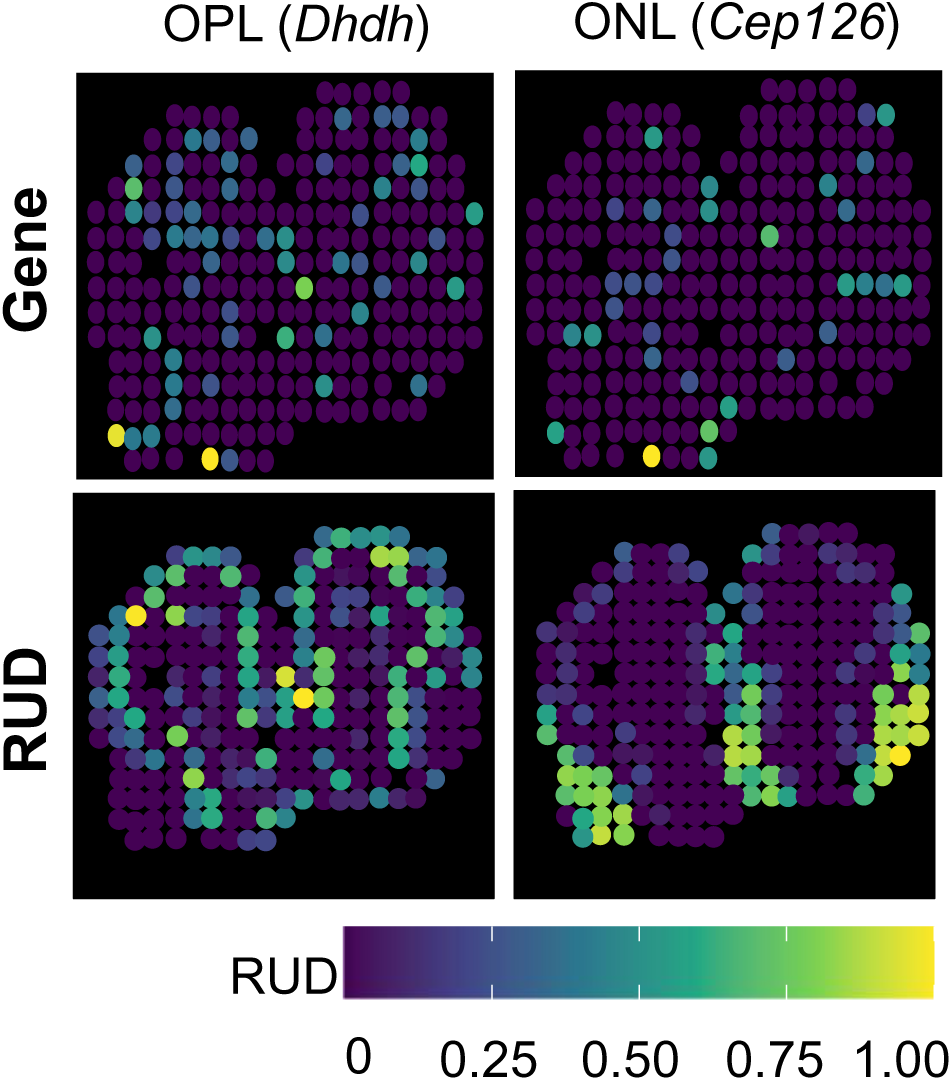
Two representative genes showing clear spatial APA usage patterns but no gene expression patterns. The ‘Gene’ row is the average gene expression level of the respective gene in all spots which represented by the sum of expression levels of poly(A) sites in the gene. The ‘RUD’ row is the average RUD scores of the gene in all spots.

**Figure S4.**
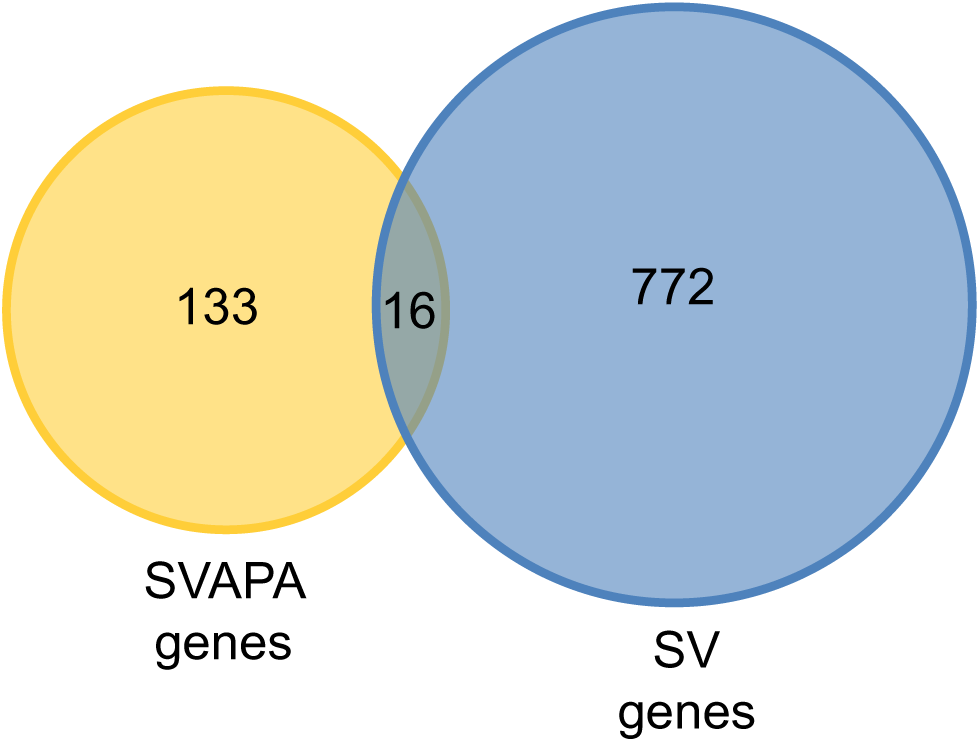
Overlap between spatially variable APA (SVAPA) genes and SV genes. The 16 overlap genes are *Sft2d2, Runx1t1, Epha5, Gabrb2, Gja1, Igfbp4, Tmem132b, Fam149a, Camk1d, Ak5, Pde5a, Rbfox1, Crtc1, Grasp, Magt1 and Fibcd1*.

**Figure S5.**
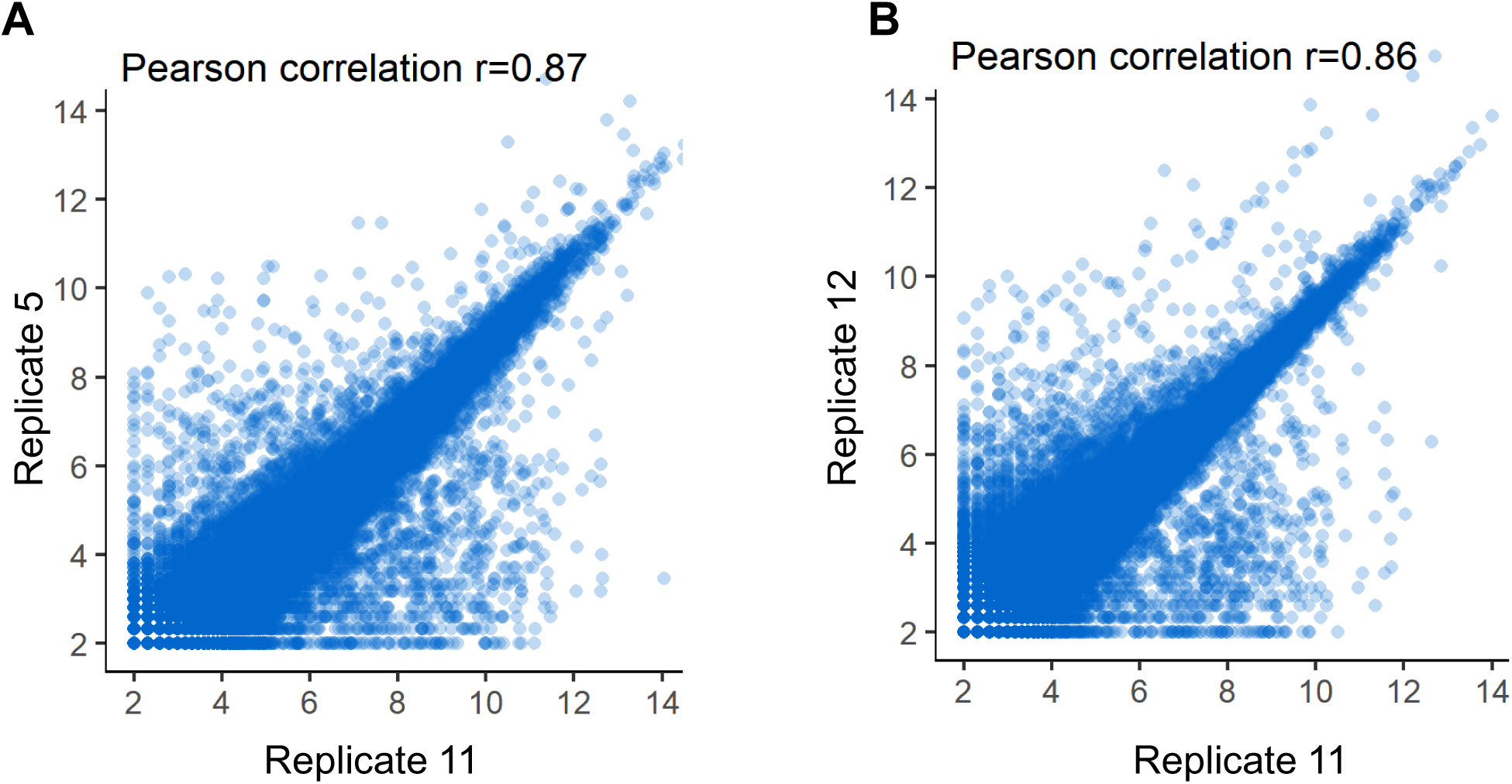
Scatter plots showing expression levels of poly(A) sites between replicates. **A.** Correlation between Replicate 5 and 11. **B.** Correlation between Replicate 5 and 12. Each dot is one poly(A) site and the axis is log2 scaled.

**Figure S6.**
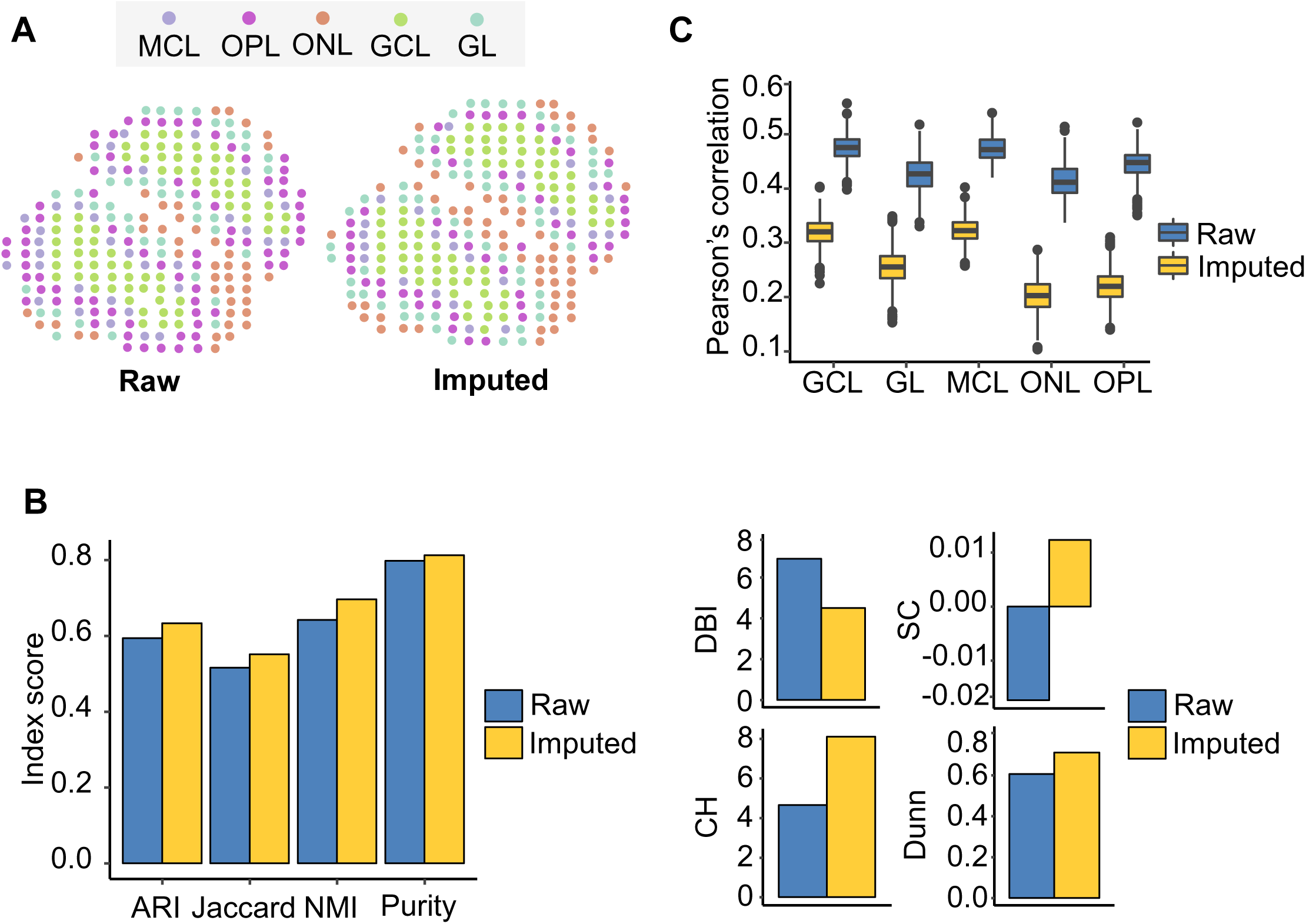
Validation of stAPAminer in imputing APA signals using Replicate 5 of ST MOB. **A.** Visualization of ST spots on the tissue image before (raw) and after (imputed) APA imputation. **B.** Evaluation of the performance of the imputation model. Four metrics were used for evaluating the performance in the context of clustering, including ARI, Jaccard, NMI, and Purity, and four internal validation metrics without relying on the reference labels were also used, including DBI, CH, SC, and Dunn. **C.** Boxplot showing Pearson’s correlations between spot pairs in each layer estimated using imputed and the raw RUD scores. For each layer, correlations of all pairwise spots were calculated.

**Figure S7.**
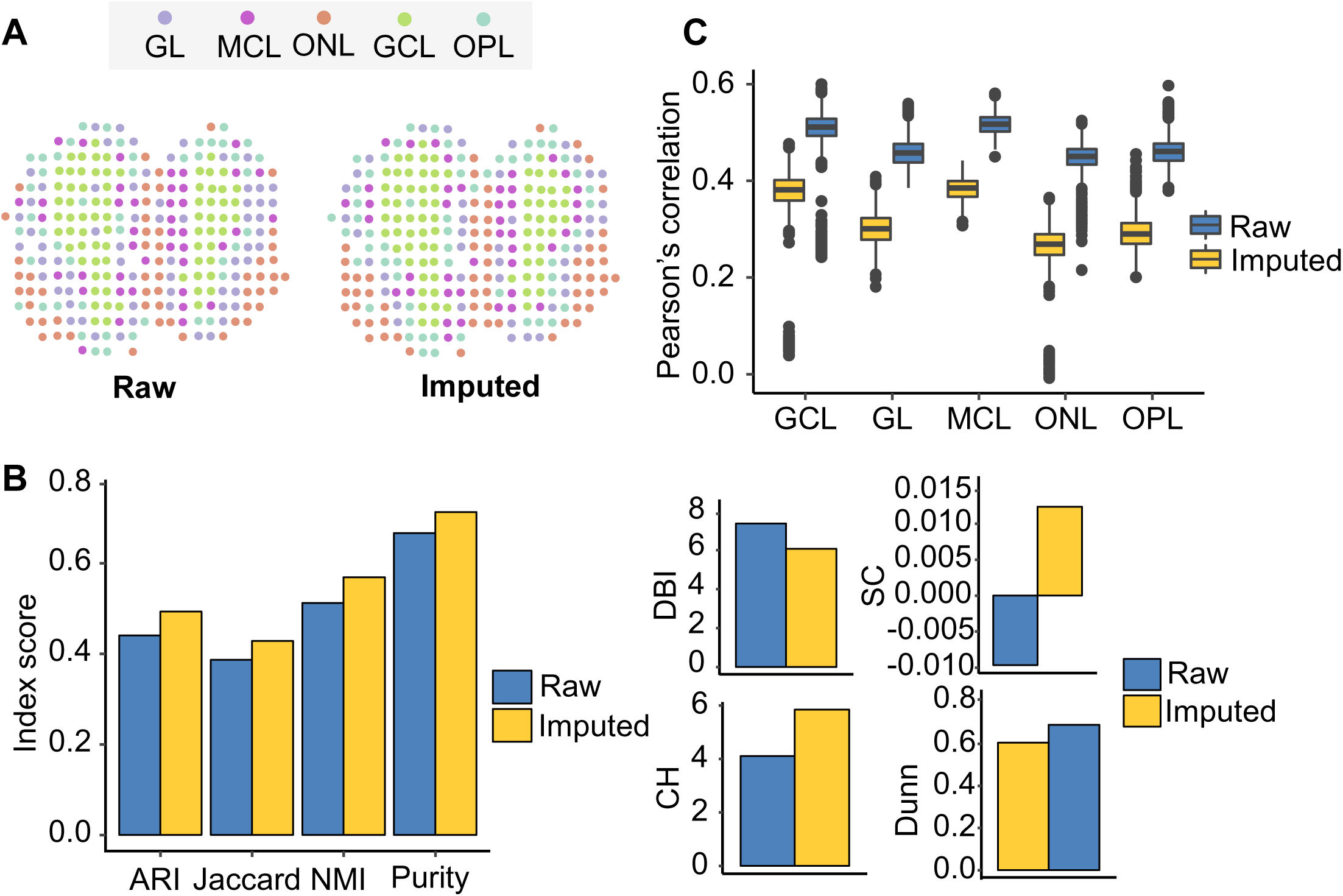
Validation of stAPAminer in imputing APA signals using Replicate 12 of ST MOB. **A.** Visualization of ST spots on the tissue image before (raw) and after (imputed) APA imputation. **B.** Evaluation of the performance of the imputation model. Four metrics were used for evaluating the performance in the context of clustering, including ARI, Jaccard, NMI, and Purity, and four internal validation metrics without relying on the reference labels were also used, including DBI, CH, SC, and Dunn. **C.** Boxplot showing Pearson’s correlations between spot pairs in each layer estimated using imputed and the raw RUD scores. For each layer, correlations of all pairwise spots were calculated.

**Figure S8.**
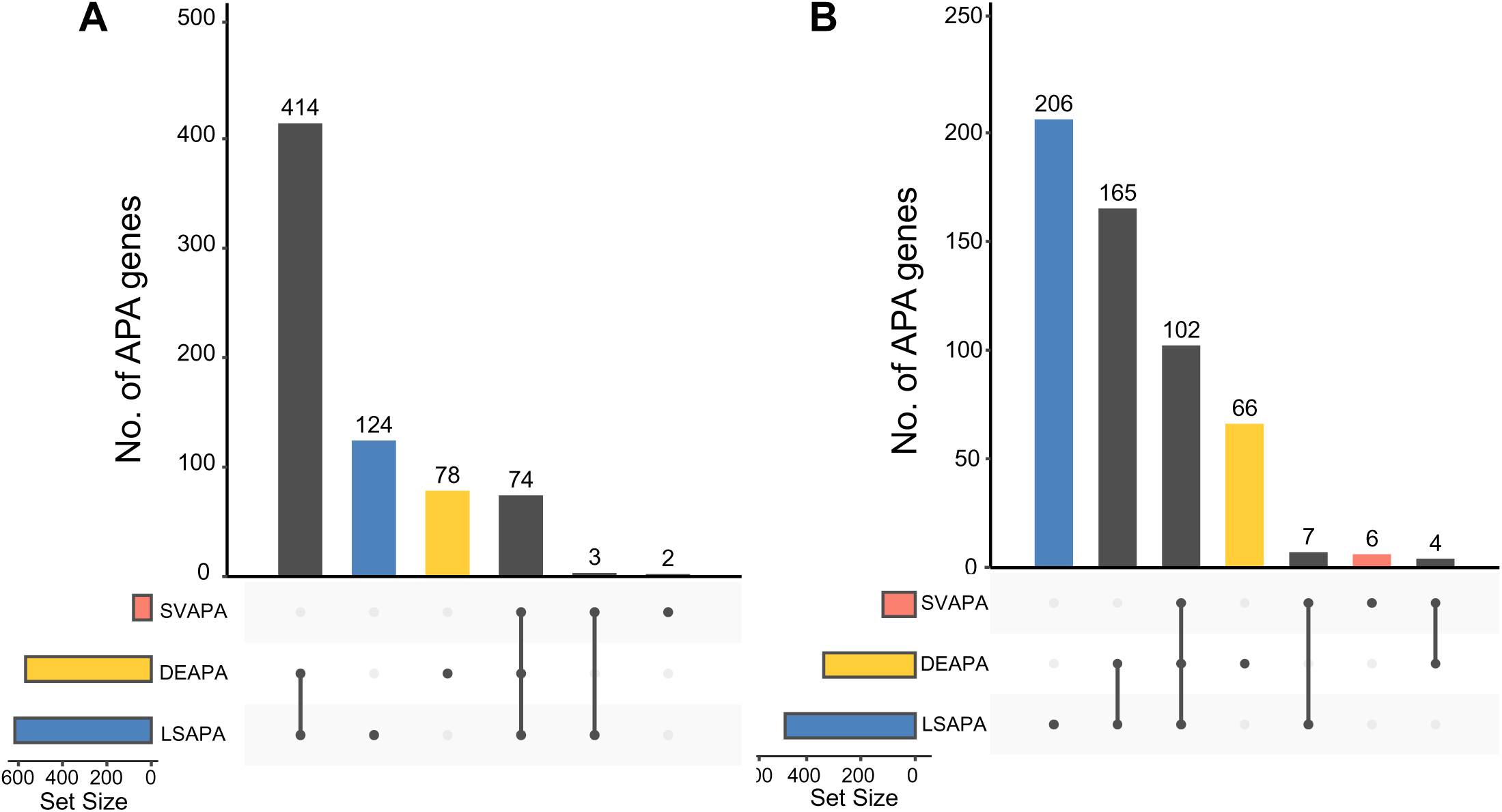
Upset plot showing overlap of DEAPA, LSAPA, and SVAPA genes. **A.** The result for Replicate 5. **B.** The result for Replicate 12.

**Figure S9.**
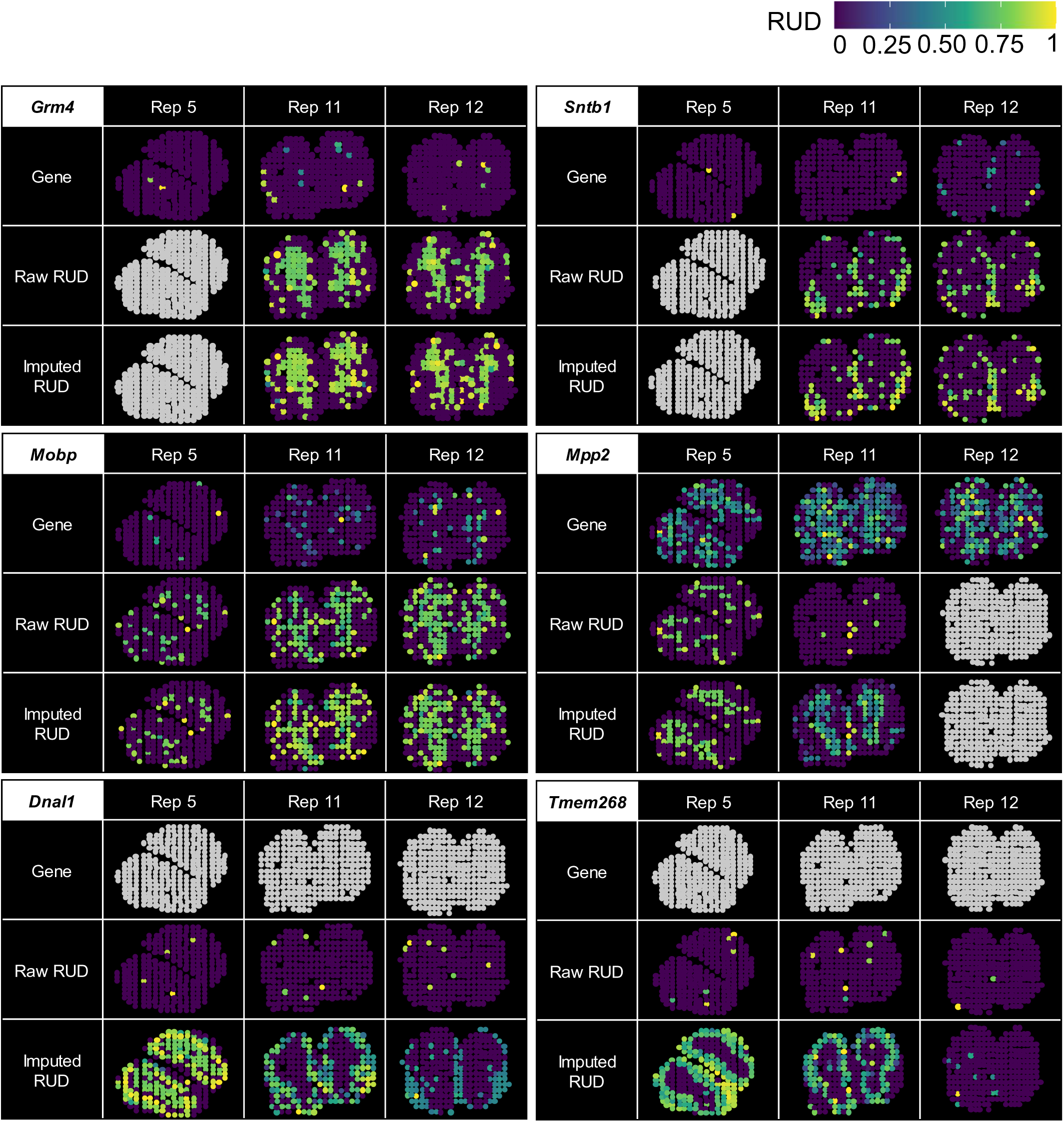
Representative genes with spatially variable APA usages detected from the three replicates of ST MOB. Tissue images of the respective gene based on the scaled gene expression level (top), raw RUD scores (middle) and imputed (bottom) RUD matrices were shown. Color represents RUD scores or scaled gene expression levels (yellow, high; blue, low).

**Table S1 Poly(A) sites identified from ST-MOB using scAPAtrap**

**Table S2 DEAPA genes between each pair of morphological layers**

**Table S3 Differentially expressed genes identified using the gene-spot expression matrix**

**Table S4 List of representative genes in the olfactory system presented in previous studies or public resources**

**Table S5 LSAPA genes by comparing the RUD profile of a layer to all other layers**

**Table S6 SVAPA genes identified by SPARK**

**Table S7 SV genes identified by SPARK**

**Table S8 Gene ontology (GO) terms enriched in the combined list of DEAPA, LSAPA, and SVAPA genes**

**Table S9 DEAPA, LSAPA, and SVAPA genes identified for Replicate 5 of ST-MOB**

**Table S10 DEAPA, LSAPA, and SVAPA genes identified for Replicate 12 of ST MOB**

**Table S11 DEAPA genes of intronic APA between each pair of morphological layers**

**Table S12 LSAPA genes of intronic APA by comparing the intronic ratio profile of a layer to all other layers**

**Table S13 SVAPA genes that display spatial intronic APA patterns identified by SPARK**

**Table S14 Top 10 GO terms enriched in the combined gene list of DEAPA, LSAPA, and SVAPA genes with intronic APA**

